# Nanodisc-Mediated Visualization of Crowding-Induced Condensation and Membrane Reorganization in Two-Dimensional Membrane Environments

**DOI:** 10.64898/2026.01.02.697448

**Authors:** Ru-Hsuan Bai, Hsin-Yi Tien, Chu-Chun Cheng, Hong-Jou Tsai, Che-Kai Lin, Chin-Ju Kuo, Wei-Tsung Wong, Chang-Ming Jiang, Yun-Wei Chiang, Chun-Wei Lin

## Abstract

Although biological membranes exhibit complex and dynamic organization, the mechanistic role of molecular crowding in governing lateral membrane heterogeneity remains poorly characterized experimentally. Here, we provide direct experimental visualization showing that crowding-induced condensation of membrane-anchored macromolecules above the bilayer interface reorganizes membrane dynamics and gives rise to spatially heterogeneous lipid mobility. Both PEGylated lipids and membrane-anchored proteins undergo surface-density-driven condensation on supported lipid bilayers, forming immobile regions that constrain lipid diffusion. Notably, the condensation threshold varies inversely with PEG chain length and surface density, defining a quantitative relationship between molecular size and crowding strength. To directly visualize these crowding-induced structures, we employed nanodelivery using lipid-loaded nanodiscs, revealing a clear correspondence between condensed crowder regions and diffusion barriers reminiscent of the picket-fence model of live-cell membranes. Similar condensation behavior observed for protein-crowded SLBs demonstrates the generality of this crowding-driven mechanism. Together, these findings establish surface-density-driven crowding and condensation of membrane-anchored macromolecules as a key physical mechanism underlying lateral membrane inhomogeneity and position nanodelivery as a general approach for interrogating membrane organization across synthetic and biological systems.

## Introduction

The cellular interior is an extraordinarily crowded and heterogeneous environment, with macromolecules occupying roughly 35% of the total volume (1, 2). In such a confined space, molecules cannot overlap, leading to steric exclusion and repulsive interactions. This phenomenon, first described by Minton in 1983 as the excluded-volume effect (3), arises from neighboring molecules that limit accessible space, restrict conformational freedom, and reduce diffusion rates, while effectively increasing local concentrations (4). These crowding-induced consequences have profound impacts on cellular biochemistry. For example, molecular crowding has been shown to promote protein folding (5–7), alter conformational stability (8, 9), accelerate association reactions (10–12), and drive phase separation in aqueous solutions (13–16). Beyond these effects, crowding is increasingly recognized as an active regulatory mechanism, with cells tuning crowding levels to modulate biochemical processes (17).

In addition to the cytosol, the plasma membrane is a highly heterogeneous and dynamic system, composed of diverse lipids and abundant membrane proteins often decorated with glycans (18–20). The recruitment of cytoplasmic proteins to the membrane surface further amplifies crowding, making it an intrinsic property of cellular membranes. Confinement of biochemical processes to the two-dimensional membrane environment has been shown to substantially alter reaction kinetics and regulatory behavior, underscoring the importance of membrane organization in dictating biological function (21). Crowding shapes membrane physical characteristics and modulates interfacial processes. For instance, large crowding agents can exclude macromolecules from the bilayer, generating osmotic imbalances that perturb membrane integrity and promote fusion (22, 23). After electrofusion, vesicles containing crowders undergo morphological transitions driven by depletion effects (24). Similarly, high densities of membrane-associated proteins can induce pronounced curvature (25–28). Together, these studies establish that molecular crowding strongly influences the architecture, dynamics, and stability of cellular membranes.

The heterogeneous organization of biological membranes has long been recognized as a hallmark of their structure and function (29–33). In 1978, lipids were shown to segregate into distinct domains characterized as either liquid-disordered (Ld) or liquid-ordered (Lo) phases (34). During the following decade, researchers reported that membrane proteins can undergo condensation concomitant with lipid phase transitions (35–38). By the 1990s, attention shifted to the role of cholesterol in modulating lateral heterogeneity (39, 40). In 1997, the concept of lipid rafts—ordered membrane domains enriched in sphingomyelin and cholesterol—was introduced (41, 42). More recently, thermally driven phase separation has been shown to arise from differences in lipid molecular structures and packing properties (43). Beyond lipid phase behavior, the picket-fence model was proposed to explain the anomalous hop diffusion of lipids and proteins in the plasma membrane (44–46). In this framework, the membrane is supported by an underlying actin cytoskeleton, where transmembrane actin-binding proteins serve as “pickets” that restrict diffusion, creating nanoscale compartments. These barriers give rise to relatively immobilized nanoclusters of membrane proteins (47). This model provides a structural basis for understanding how the three-dimensional organization of the plasma membrane emerges from the interplay between cytoplasmic and extracellular components (47, 48). While lipid composition, cholesterol, and cytoskeletal interactions are known to govern membrane heterogeneity, whether molecular crowding can itself drive phase transitions within the membrane plane has not been experimentally clarified, largely because direct molecular-level visualization of crowding-induced organization has remained technically challenging.

Here, we investigate how molecular crowding modulates membrane organization and dynamics using supported lipid bilayers (SLBs) as a tractable model. Poly(ethylene glycol) (PEG)–conjugated lipids were incorporated as a well-established molecular crowder (11, 49, 50). Embedding PEG directly into the bilayer created a controlled crowding environment in the two-dimensional membrane plane. Previous studies suggest that PEG chains can undergo local condensation, modulating accessible free volume and enhancing system entropy (51, 52). We show that crowding of membrane-anchored PEG induces a two-dimensional phase transition of the crowders themselves, which in turn generates lateral membrane heterogeneity by partitioning the bilayer into distinct regions that impose barrier-limited diffusion. This crowding-induced pattern resembles the meshwork of the picket-fence model, establishing a direct connection between crowding-driven membrane phase transitions and cytoskeleton-dependent compartmentalization (44–46, 48).

To directly visualize PEG-driven condensation domains, we employed nanodiscs—disk-shaped lipid bilayers stabilized by membrane scaffold proteins—that enable nanodelivery of fluorescent lipids into diverse membrane systems (53–55). This adapted nanodisc-mediated approach enables direct visualization of crowding-driven condensation and molecular reorganization on membranes, revealing phenomena previously inaccessible at this level of spatial resolution. Nanodiscs have emerged as versatile tools for delivering lipids, reconstituting proteins, and altering membrane composition in cells (56–61). Under PEG-induced crowding conditions, nanodelivery revealed striking correspondence between condensate patterns and restricted lipid diffusion. In this limited sense, the observed barrier-like features bear a loose conceptual resemblance to the picket-fence model, in that both involve crowding-induced constraints on lateral diffusion. The results from our *in vitro* reconstituted system demonstrate crowding-induced compartmentalization in a simplified membrane setting, providing a physically grounded model for exploring how crowding-mediated constraints on diffusion could contribute to membrane heterogeneity in cellular contexts.

In addition to PEGylated lipids, we found that membrane-anchored fluorescent proteins on SLBs also undergo crowding-driven phase transitions. As surface density increased, proteins spontaneously assembled into immobile condensates, while the underlying bilayer remained fluid. In both cases, a phase transition of membrane-anchored crowders gives rise to lateral membrane heterogeneity as an emergent consequence, rather than the bilayer itself undergoing a classical two-dimensional phase separation. These results demonstrate that crowding from either PEG polymers or membrane-anchored proteins can serve as a general physical mechanism for generating lateral membrane heterogeneity. This in-vitro reconstituted system provides mechanistic insight into how crowding dictates membrane organization and establishes a broadly applicable model for dissecting complex membrane interactions. Because PEGylated lipids are widely employed in lipid nanoparticle (LNP) formulations for mRNA vaccines (62–66), these findings further underscore the biomedical significance of crowding-induced membrane transitions.

## Results

To investigate how molecular crowding influences lateral membrane dynamics, supported lipid bilayers (SLBs) were prepared with varying fractions of PEGylated phospholipid (PEG5000-PE). To monitor lipid mobility, 0.5 mol% FITC-DHPE was incorporated as a fluorescent probe, and fluorescence recovery after photobleaching (FRAP) was used to study lipid diffusion. Given the larger excluded-volume effect of PEG (5, 67), SLBs containing 0–1.5 mol% PEG5000-PE were analyzed (Figure 1A).

**Figure 1.**
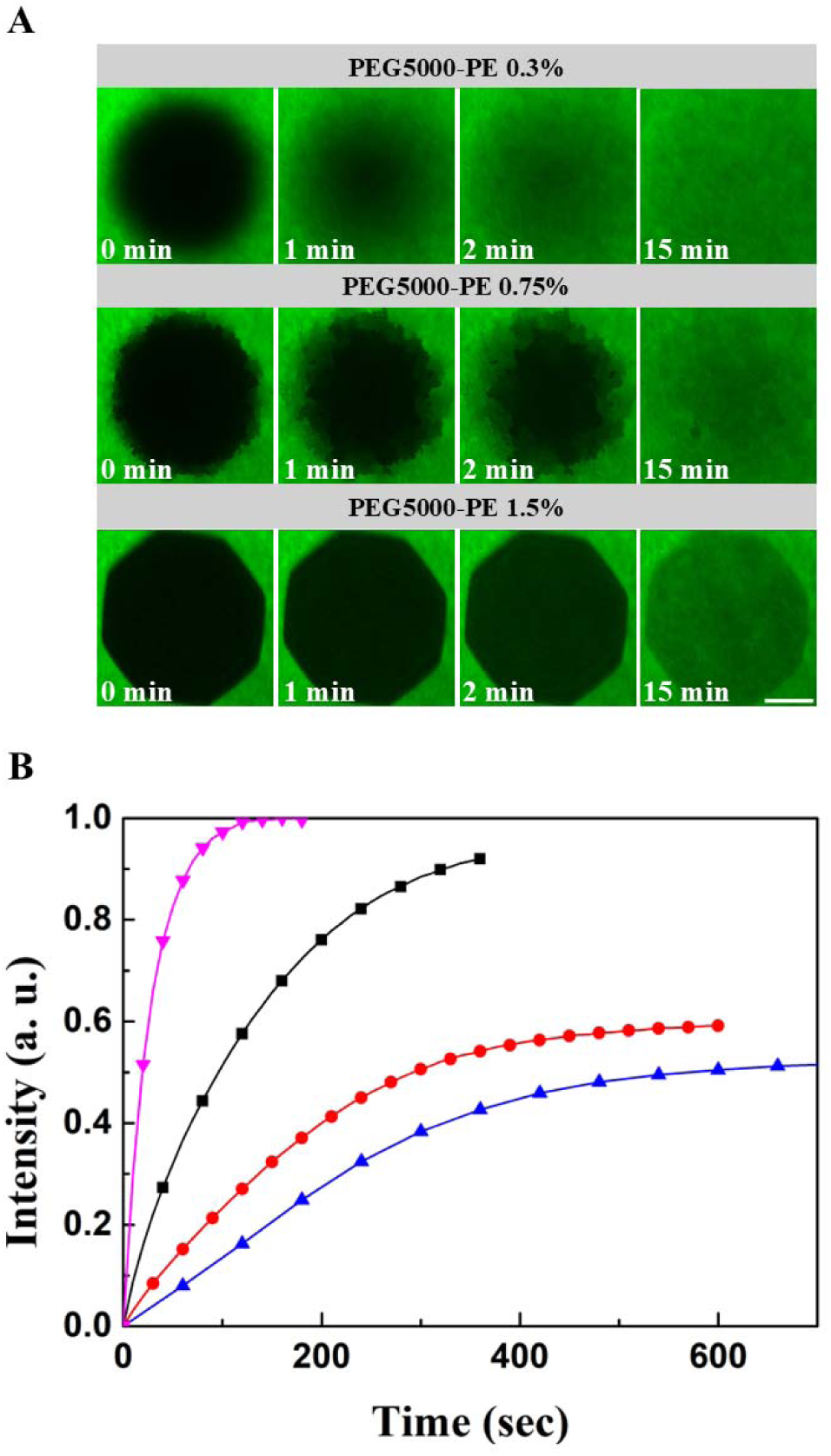
FRAP analysis of SLBs containing different fractions of PEG5000-PE. (A) SLBs composed of 0.05 mol % FITC-DHPE and 0.3–1.5 mol % PEG5000-PE were photobleached by high-power laser irradiation for 10 s, and fluorescence images were collected every 30 s to monitor recovery. The SLB with 0.3 mol % PEG5000-PE showed nearly complete fluorescence recovery (∼100 %) within 3 min. At 0.75 mol %, recovery was partial, indicating barrier-limited diffusion. At 1.5 mol %, fluorescence remained low even after 15 min, suggesting a largely immobile bilayer. Scale bar: 30 μm. (B) Fluorescence recovery curves corresponding to (A). Traces represent SLBs containing different PEG5000-PE fractions (pink inverted triangles, 0 %; black squares, 0.3 %; red circles, 0.75 %; blue triangles, 1.5 %).

At high PEG5000-PE content (1.5 mol%), lipid diffusion was markedly reduced, and fluorescence recovery remained incomplete, indicating the presence of an immobile fraction associated with the glass-supported bilayer. FRAP recovery traces further revealed that lipid mobility was progressively slowed when the PEG5000-PE fraction exceeded 0.3 mol%. Compared to crowding in three-dimensional solution, the threshold concentration required to impede two-dimensional diffusion was remarkably low (68), consistent with the enhanced local effective concentration of crowders confined to a 2D membrane (69–71).

At intermediate PEG5000-PE fractions (0.75 mol%), the average rate of fluorescence recovery alone did not capture the full behavior of the system. Instead, FRAP revealed pronounced spatial heterogeneity in the recovery profile across the photobleached region. Unlike the uniform recovery observed at PEG5000-PE fractions below 0.3 mol%, membranes containing 0.75 mol% PEG5000-PE exhibited spatially inhomogeneous, mosaic-like recovery patterns (Figure 1A), indicative of barrier-limited diffusion reminiscent of the picket-fence model of membrane compartmentalization. Importantly, this behavior reflects spatial variations in lipid mobility rather than a global slowdown of diffusion across the membrane.

Initially, this barrier-limited recovery was attributed to a potential contribution from PE headgroups. However, control experiments using comparable or higher fractions of non-PEGylated PE lipids showed no such spatially heterogeneous recovery (Figure S6), demonstrating that the PEG5000 headgroup itself is the critical determinant underlying the observed lateral inhomogeneity.

Notably, the spatially heterogeneous FRAP recovery observed here differs from classical lipid phase separation into coexisting membrane domains. Rather than reflecting a simple change in the overall diffusion rate, these results suggest spatially inhomogeneous constraints on lipid mobility imposed by membrane-anchored crowders, raising the question of how such crowders generate lateral inhomogeneity in supported lipid bilayers. To address this, we sought to directly visualize the spatial distribution of PEGylated lipids within supported lipid bilayers (SLBs).

We first reduced the fraction of the fluorescent lipid Texas Red-DHPE (TR-DHPE) to test whether membrane inhomogeneity could be revealed by uneven reporter distribution (Figure S1). At 0.75% PEG5000, immobile features were faintly detectable when TR-DHPE was lowered to 0.0006 mol%; however, these patterns did not correspond with the barrier-limited FRAP signatures in Figure 1A.

We next turned to nanodisc-mediated lipid delivery, which enables precise unloading of fluorescent lipids into bilayers (54, 55, 72–74). Nanodiscs containing 4 mol% TR-DHPE were added to SLBs from solution. This approach enabled incorporation of fluorescent lipid beyond the single-molecule regime, providing sufficient fluorescence signal to clearly distinguish TR-DHPE distribution between mobile and immobile regions of the membrane. Strikingly, nanodelivery precisely highlighted barriers formed by PEG lipid headgroups (Figure 2).

**Figure 2.**
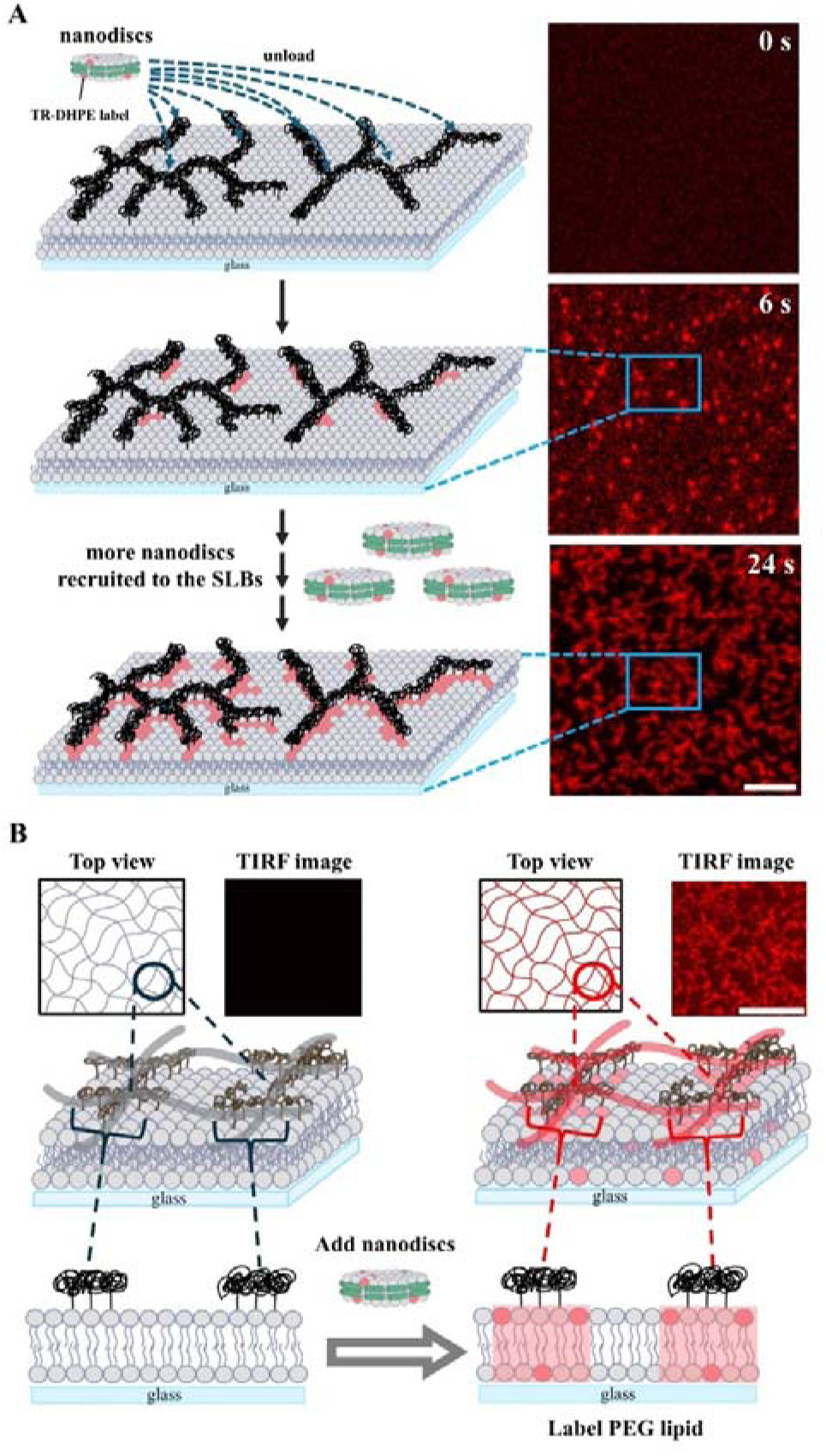
The phase transition of PEG above the SLB is detected using the nanodelivery method previously developed (72). (A) Time-lapse images show that lipids from nanodiscs are unloaded onto the SLB containing 0.75 % PEG5000-PE within 30 s. The nanodelivery-mediated lipid transfer preferentially occurs at PEG-rich condensates and the corresponding lipid domains. Texas Red–DHPE lipids, initially incorporated into nanodiscs, appear as bright fluorescent spots on the SLB. Lipids of nanodiscs are locally transferred to immobile regions of the bilayer when nanodiscs collide with PEG condensates above the SLB, corresponding to the bright fluorescence observed at t = 6 s after nanodisc addition. As more nanodiscs interact with PEGylated lipids and release Texas Red–DHPE, the accumulating signal delineates the PEG-induced phase transition on the SLB. Scale bar = 5 μm. (B) Schematic illustration showing that the PEG phase transition above the SLB drives a corresponding phase transition through the lipid anchor within the bilayer, as observed in the fluorescence images. Scale bar = 8 μm. The nanodelivery-highlighted regions are illustrated on the right side of the schematic.

From FRAP experiments (Figure 3A and 3C), these barriers were immobile: photobleached regions recovered to only ∼20% of their initial intensity after 3 min, indicating limited lipid exchange between barrier structures and the surrounding bilayer. Nanodisc-delivered fluorescent lipids also revealed PEG-driven barriers whose static nature is consistent with the FRAP behavior of SLBs containing 0.75% PEG5000.

**Figure 3.**
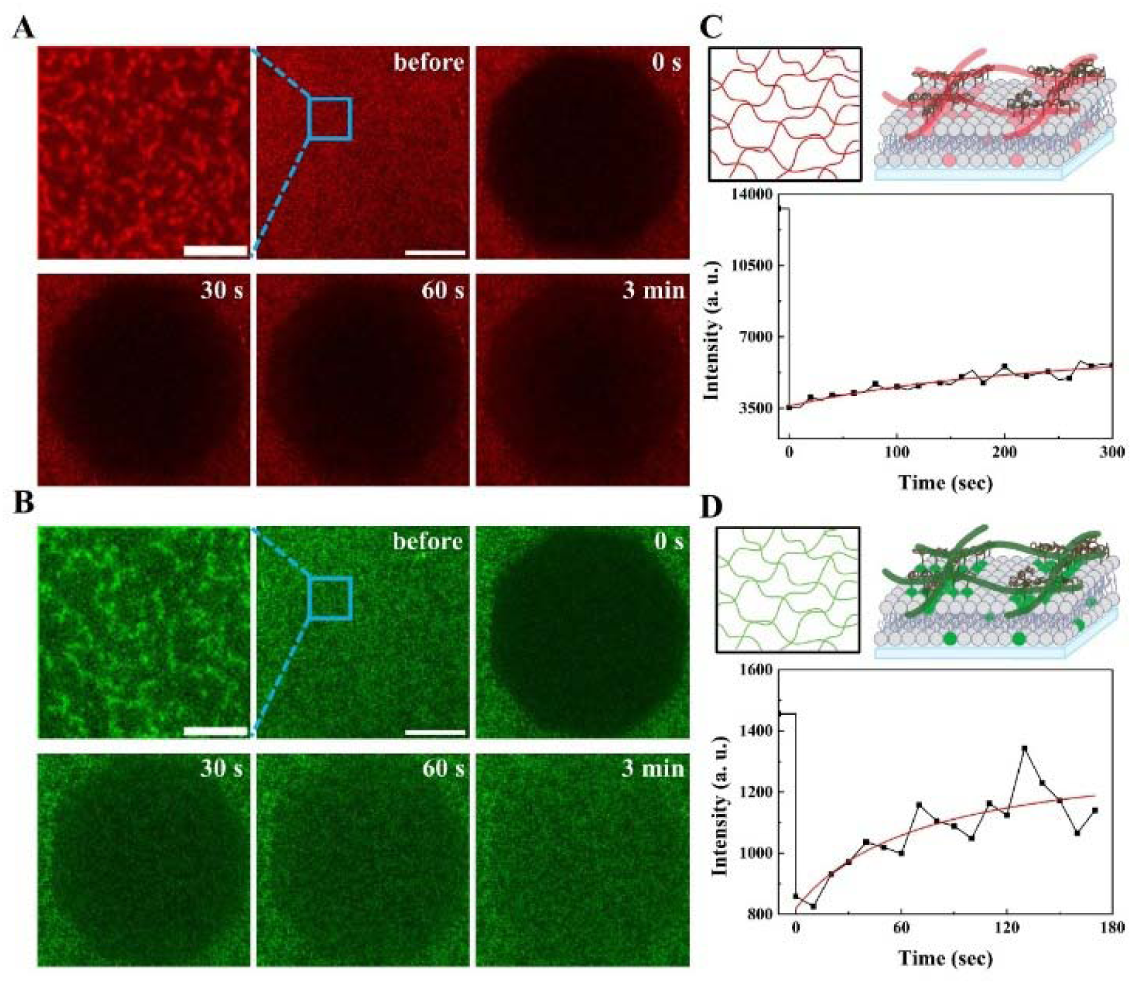
The phase separation of the SLB driven by the crowding-induced phase transition of PEG above the membrane, visualized using nanodiscs loaded with different fluorescent lipids. (A) The SLB containing 0.75 % PEG5000-PE undergoes phase separation, which is highlighted by nanodiscs carrying Texas Red–DHPE (TR-DHPE). FRAP measurements performed on this SLB reveal partial fluorescence recovery, indicating restricted lipid mobility within the immobile domains. (B) A similar phase separation is observed when nanodiscs containing FITC-DHPE are used. In this case, FRAP shows faster fluorescence recovery in both the mobile and immobile regions, reflecting differences in local membrane dynamics between these two chromophores. (Scale bars: time-lapse = 5 µm; FRAP = 25 µm.)

After visualizing barriers on SLBs using nanodelivery, we investigated how fluorescent lipids report on these structures. TIRF microscopy captured the initial unloading of TR-DHPE lipids from nanodiscs into the SLB (Figure 2B). In control experiments without barriers, lipids dispersed homogeneously across the membrane (Figure S2). By contrast, when PEGylated lipids had already induced barrier formation, TR-DHPE preferentially accumulated at these regions. Quantitative analysis showed that fluorescence intensity increased more rapidly at the barrier than in the surrounding regions (Figure S3), indicating preferential partitioning of TR-DHPE into barrier domains (Figure 2). When the SLB was not overloaded with fluorescent lipid, nanodisc-delivered lipids still effectively enabled barrier labeling with fluorescent lipids, consistent with the schematic models in Figures 2 and 3.

To confirm that barrier labeling with fluorescent lipids was not dye-specific, nanodiscs loaded with FITC-DHPE were also employed, yielding comparable visualization of the PEG-induced barriers (Figure 3B). Both FRAP measurements in Figure 3 revealed recovery behavior associated with the crowding-induced condensed regions induced by PEG, consistent with lipid exchange between the condensed barriers and the surrounding bilayer while maintaining reduced mobility in these barrier regions. Notably, the morphology of PEG5000-PE–induced barriers did not resemble canonical binodal or spinodal phase separations, but instead reflected crowding-driven lipid organization consistent with PEG-induced phase transitions illustrated schematically in Figures 2 and 3.

To explore how the molecular size of the PEG headgroup influences the phase transition on supported lipid bilayers (SLBs), lipids with different PEG chain lengths were incorporated into the bilayer, and nanodiscs were introduced above the SLB to capture the extent of phase separation (see Figures 4 and S5). Similar to PEG5000-PE, other PEGylated lipids also induced phase transitions at characteristic mole percentages. These experiments revealed that each PEG chain length exhibited a distinct threshold concentration for inducing phase separation. Control SLBs containing an equivalent percentage of PE lipids without the PEG modification showed no detectable phase transition, confirming that the observed transitions arise specifically from PEG headgroups rather than from lipids alone (see Figure S6). Nanodelivery labeling showed that the morphological appearance of these transitions ranged from isolated clusters to mesh-like networks, and ultimately to nearly uniform layers when both PEG content and chain length were high. Across the five PEG-lipid species tested, larger PEG headgroups drove phase transitions at lower mole percentages, indicating that the effective size of the crowding moiety directly governs the onset of PEG-induced phase separation on SLBs.

**Figure 4.**
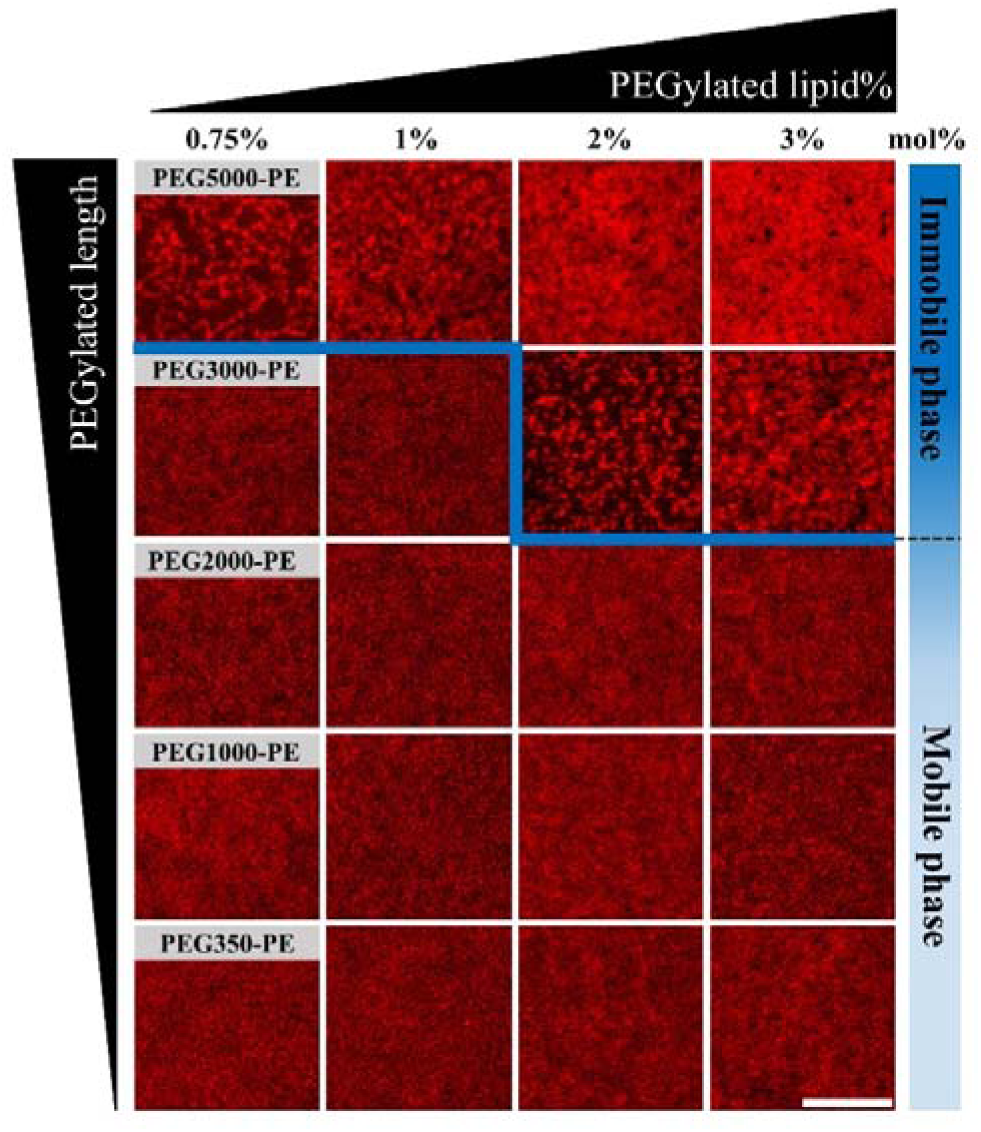
The phase transition of the SLB is mediated by the mole percentage of PEG lipid (0.75–3 %) and by the length of the PEG chain, using PEG5000-PE, PEG3000-PE, PEG2000-PE, PEG1000-PE, and PEG350-PE from the top row to the bottom row at 25 °C. The complete dataset is shown in Figure S5. (Scale bar = 8 µm.) Once the membrane undergoes phase transition driven by the crowding effect, the fluorescence image exhibits inhomogeneity, with the immobile phase preferentially labeled by fluorescent lipids delivered from nanodiscs. The area of the immobile phase increases progressively as the PEG-lipid percentage increases. PEG lipids with longer polymer chains enable phase transition at lower PEG-lipid fractions, indicating that the crowding threshold depends on the effective molecular size of the tethered PEG group.

Given that the observed phase transition did not resemble canonical binodal or spinodal demixing, we next examined the physical properties of the crowding-induced membrane inhomogeneity. Two complementary fluorescent lipids, FITC-DHPE and TR-DHPE, were employed to probe distinct aspects of the supported lipid bilayer (SLB). FITC-DHPE (0.05 mol %) was incorporated directly into the bilayer to monitor membrane dynamics by FRAP, whereas TR-DHPE was delivered via nanodiscs to selectively label crowding-induced barriers (Figure 5).

**Figure 5.**
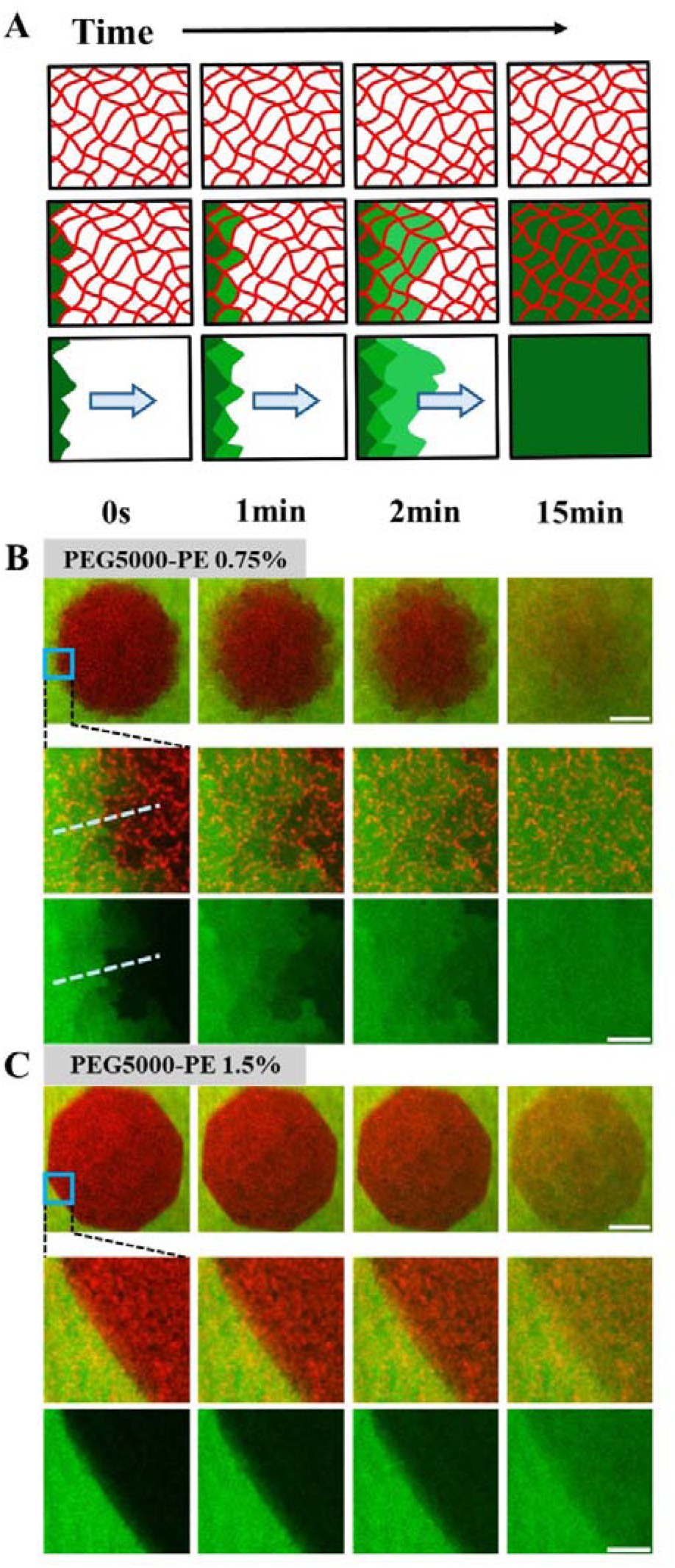
The recovery of the fluorescence signal is dictated by the crowding-induced phase transition on the SLB observed in the FRAP experiments. (A) The cartoon schematic illustrates the stepwise fluorescence recovery pattern corresponding to the framework of immobile regions formed during the phase transition driven by the crowding effect. (B) The FRAP experiment was performed on an SLB containing 0.5 → 0.05 mol % FITC-DHPE and 0.75 mol % PEG5000-PE lipids (green). The phase transition associated with PEG5000-PE is highlighted by nanodelivery labeling (red). The recovery of FITC-DHPE fluorescence closely followed the spatial pattern of the PEG-driven phase transition, leading to a characteristic stepwise increase in fluorescence intensity (see Figure S7), corresponding to the line-scan profiles in the first images of the second and third rows. (C) A similar FRAP experiment was performed on an SLB containing a higher percentage (1.5 mol %) of PEG5000-PE. Due to the more densely packed immobile phase arising from the greater PEG5000 content, fluorescence recovery was substantially slower and lacked the stepwise intensity increase observed in panel B. The agreement between the spatial distribution of the immobile phase and the recovery pattern in the mobile phase accounts for the distinct fluorescence recovery behavior initially observed in Figure 1. (Scale bars: zoom-out, 25 μm; zoom-in, 5 μm.)

By overlaying FRAP images of FITC-DHPE with TR-DHPE fluorescence images, we found that regions of restricted recovery aligned precisely with TR-DHPE–labeled barriers, indicating that PEGylated lipids generate spatially confined diffusion domains. Quantitative correlation between FRAP recovery and TR-DHPE intensity confirmed this relationship (Figure S7). The fluorescence intensity profile of TR-DHPE along the dash line in Figure 5B corresponded directly to the stepwise recovery observed by FRAP using FITC-DHPE as the probe.

Furthermore, FRAP recovery was progressively suppressed at higher PEG-lipid content. At 1.5 mol % PEG5000-PE, the barrier density was markedly increased, reducing recovery to only ∼40% after photobleaching (Figures 1 and 5C). This slowdown in fluorescence recovery indicates that densely packed PEG domains result in highly effective diffusion barriers through the associated lipid parts, substantially limiting lateral lipid mobility across the bilayer.

The crowding-induced inhomogeneity was next examined at the single-molecule level (Figure S8). Alexa Fluor 647–labeled TEV protease, anchored to the bilayer through His-tag chemistry in the presence of 0.75 mol % PEG5000-PE, exhibited constrained two-dimensional Brownian trajectories confined within TR-DHPE–marked domains. These crowding-induced barriers effectively partition the membrane into discrete compartments, closely resembling the picket-fence model of the plasma membrane in live cells (44–46, 48, 75, 76).

To examine whether protein crowding alone can drive phase transitions on membranes, we systematically increased the surface density of fluorescent proteins anchored to SLBs via click chemistry (Figure 6A). At low densities, proteins were uniformly distributed and exhibited free lateral mobility, as confirmed by FRAP measurements. As the density increased, the proteins spontaneously condensed into bright, immobile domains, indicating a crowding-induced transition to an immobile, condensed state. The transition was observed for multiple proteins, including mScarlet, mCherry, mNeonGreen, and eGFP, indicating that this behavior is not specific to a single protein but reflects a general crowding-driven phenomenon. Notably, the membrane itself remained intact and fluid, as verified by lipid FRAP controls, demonstrating that the observed inhomogeneity arose specifically from the protein layer rather than from bilayer disruption (Figure S9–S11).

**Figure 6.**
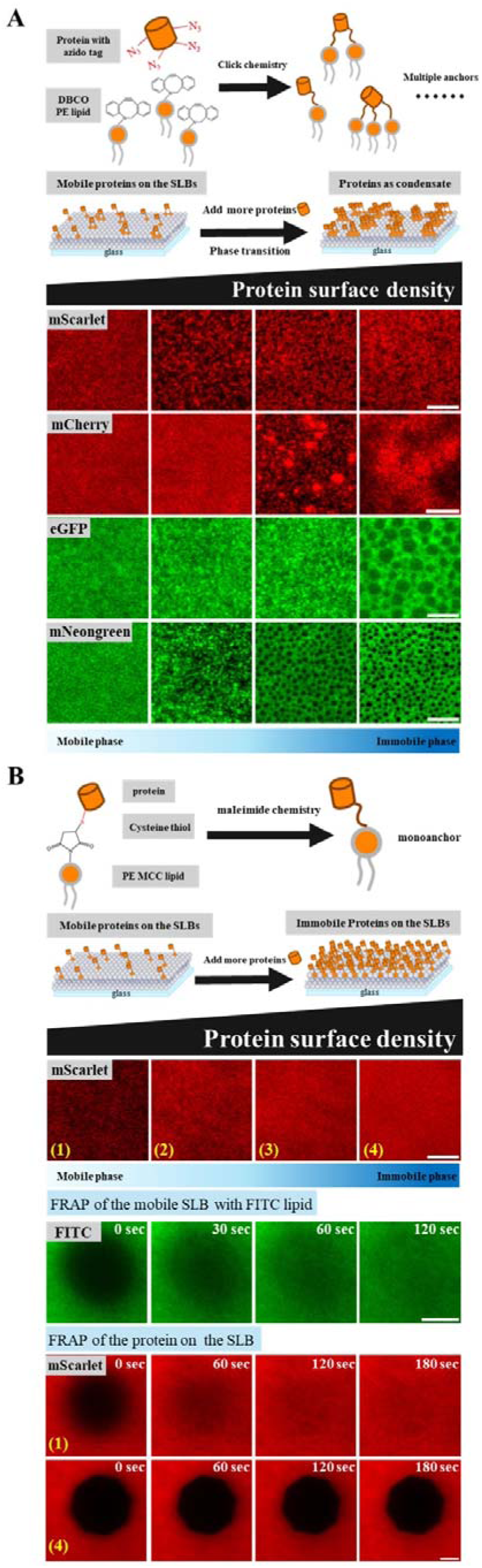
The phase transition of proteins on the SLB is driven by surface crowding conditions. (A) Different fluorescent proteins, including mScarlet, mNeonGreen, and eGFP, are anchored on the SLB via click chemistry and undergo crowding-induced phase transition as their surface density increases from approximately 100 to 1800 molecules μm[², resulting in bright, immobile protein condensates on the SLB. The cartoon schematic illustrates that the fluorescent proteins undergo phase transition when their surface density is high. Multiple membrane anchors of the proteins may further facilitate condensation on the SLB. The fluorescence images at the bottom show the transition from a homogeneous and mobile distribution across the membrane to immobile protein condensates. (Scale bar = 5 μm.) The corresponding FRAP experiments for both the membrane and the fluorescent proteins are shown in Figures S9–S12, demonstrating that the membrane remains intact while inhomogeneity arises specifically from the proteins. (B) Similar experiments were performed with fluorescent proteins anchored to the SLB using maleimide chemistry; together with His-tag chemistry shown in Figure S13, these approaches serve as monoanchoring strategies (single membrane attachment) for comparison with part A. The resulting protein condensates appear relatively uniform, and the fluorescent proteins exhibit negligible mobility in FRAP measurements. In contrast, FRAP of FITC-labeled lipids indicates that the SLB itself remains fluid. These observations demonstrate that surface crowding alone is sufficient to drive protein condensation on membranes, while multivalent anchoring can modulate the extent or sharpness of the condensed state.

Because the click-chemistry anchoring strategy can introduce multiple lipid attachments per protein and thus potentially facilitate multivalent interactions or crosslinking, we next examined whether similar condensation behavior could be observed using anchoring schemes that restrict proteins to a single membrane attachment. When fluorescent proteins were tethered to the SLB via monoanchoring strategies, including maleimide conjugation (Figure 6B and Figure S12) and Histag–mediated binding (Figure S13), we continued to observe a pronounced reduction in mobility and the formation of immobile protein-rich regions at elevated surface densities. Although the resulting condensates were generally less sharply defined than those observed for proteins anchored via click chemistry, their emergence under high surface-density conditions demonstrates that increased two-dimensional crowding alone is sufficient to induce protein condensation on membranes, without requiring multivalent anchoring.

Having established the crowding-induced condensation of membrane-anchored proteins, we next tested whether nanodelivery could serve as a generalizable approach to visualize these condensates. When mScarlet was anchored at high density to the SLB, nanodiscs loaded with 4% FITC-labeled lipid colocalized strongly with the immobile protein domains (Figure 7). This finding shows that nanodelivery provides a versatile and effective means to label protein condensates, paralleling its ability to mark PEG-driven crowding domains, and thereby offers a generalizable strategy for detecting crowding-induced membrane inhomogeneities.

**Figure 7.**
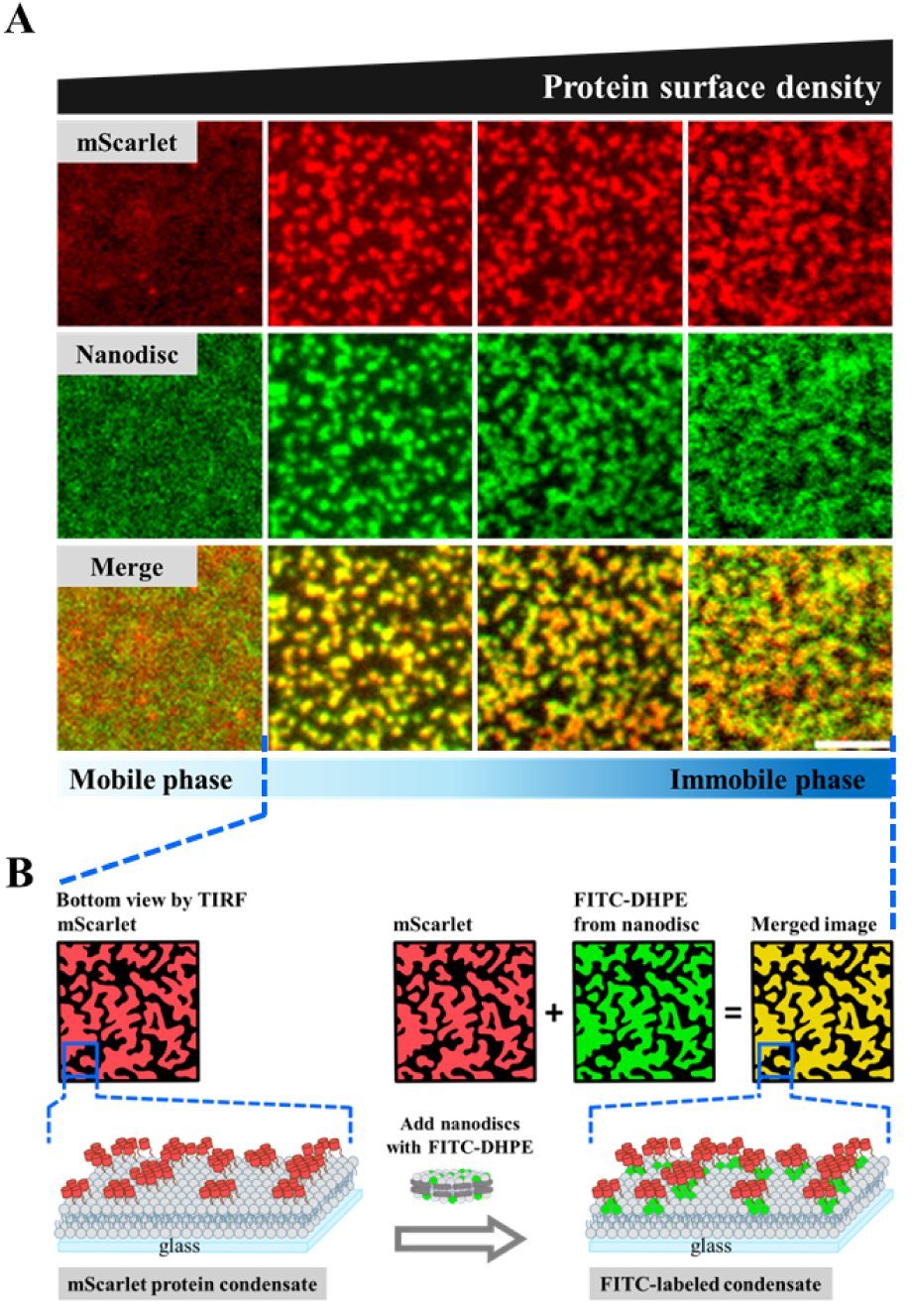
Phase transition of membrane-associated mScarlet induced by surface crowding and visualized through nanodelivery. (A) mScarlet was covalently anchored to the supported lipid bilayer via click chemistry. Increasing the surface density of mScarlet on the membrane established the crowding condition, leading to the formation of protein condensates. To probe the local membrane environment, nanodiscs composed of MSP and phospholipids doped with 4% FITC-conjugated lipid were applied to the bilayer for 10 min, followed by washing to remove free nanodiscs. The fluorescent lipid cargo was successfully unloaded and served as a secondary reporter of local membrane organization. The merged fluorescence images (mScarlet in red, FITC in green) show strong colocalization, indicating that mScarlet condensates overlap with membrane inhomogeneities revealed by nanodelivery. These inhomogeneities resemble the PEG-induced condensation domains observed in PEGylated bilayers, consistent with the barrier-limited diffusion mechanism illustrated in Figure 2. (Scale bar: 5 μm) (B) Schematic illustration of the surface-crowding-induced phase transition of membrane-anchored mScarlet and its visualization through nanodelivery.

## Discussion

### Crowding-Induced Lipid Inhomogeneity in Supported Lipid Bilayers

The dynamics of SLBs are strongly influenced by the crowding effect at the membrane surface (see Figure 1). In addition to the slower fluorescence recovery expected from lipids bearing bulky crowding moieties, our experiments reveal two distinct FRAP recovery modes. One corresponds to barrier-limited recovery, while the other exhibits only partial recovery, reflecting a substantial immobile lipid fraction within SLBs. Importantly, Figure 1 primarily reports spatially heterogeneous FRAP recovery that depends on the surface density of membrane-anchored PEG crowders. While such behavior is consistent with collective organization under crowding, Figure 1 alone does not uniquely distinguish between a thermodynamic phase transition and other crowding-induced mechanisms.

Motivated by this spatially heterogeneous recovery behavior, we next sought to investigate the origin of the observed membrane inhomogeneity. Specifically, we asked whether the observed inhomogeneity arises from lateral phase separation of SLB lipids. When a small fraction of fluorescent lipids was incorporated into SUVs during SLB preparation, the resulting bilayers displayed predominantly homogeneous fluorescence without discernible features corresponding to the inhomogeneities observed in Figure 1 (see Figure S1).

To overcome this limitation, we adopted the nanodelivery approach pioneered in our laboratory and others, which enables precise incorporation of trace amounts (down to the single-molecule level) of fluorescent lipids into SLBs. Without this method, subtle differences in lipid affinity for distinct SLB domains could not be resolved. Nanodelivery enables these distinctions to be visualized through the differential partitioning of fluorescent lipids between regions exhibiting reduced mobility and regions that remain comparatively mobile within the same bilayer.

### Condensation above the Membrane as the Mechanistic Basis for Crowding-Induced Heterogeneity

Upon formation of the SLB from SUVs on glass substrates, the crowding reagent PEG builds up above the membrane and reaches locally high concentrations, where it undergoes a crowding-driven condensation transition governed by surface density rather than bulk concentration. In this framework, increasing the surface density of membrane-anchored PEG plays a role analogous to increasing pressure in a confined chamber, driving the system across a condensation threshold. The resulting polymer condensates, protruding above the bilayer plane, are more accessible to nanodiscs in solution. This accessibility facilitates localized nanodelivery events, preferentially inserting fluorescent and other lipids into the PEG-condensed, low-mobility membrane regions. Within only a few seconds, polymer condensate–associated immobile domains become rapidly labeled with fluorescent lipids, as shown in Figure 2.

By contrast, when fluorescent lipids are incorporated into the SUV lipid mixture prior to the formation of the SLB, this preferential enrichment is lost, resulting in a uniform fluorescence distribution even in the presence of PEG lipids (Figure S1). These observations underscore nanodelivery as a uniquely powerful strategy for resolving inhomogeneity that emerges as a consequence of condensation of membrane-anchored crowders, rather than intrinsic lipid–lipid phase separation.

At high PEG5000-PE fractions, fluorescence recovery is strongly suppressed, raising the possibility that coupling between the bilayer and the solid support could contribute to the apparent immobile fraction. However, as shown below, an analogous reduction in mobility and spatially heterogeneous recovery is observed in a distinct system in which fluorescent proteins are tethered exclusively to the membrane surface and do not interact with the substrate. This comparison indicates that the dominant contribution to the observed immobility arises from condensation and crowding of macromolecules anchored to the membrane interface, rather than from substrate-induced pinning of the bilayer.

The immobile features observed in SLBs suggest that these structures arise from components beyond the intrinsic lipid composition, as SLBs containing equal or greater amounts of PE lipid exhibit uniformly distributed fluorescence (Figure S6). Thus, the observed inhomogeneity reflects a secondary reorganization of lipid mobility and spatial distribution in response to PEG condensation, rather than a primary thermodynamic phase transition of the lipid bilayer itself. Previous work by Chung et al. (77) demonstrated that membrane-associated protein phase transitions can similarly induce lipid reorganization, supporting the view that crowding-driven condensation of surface-tethered species provides a general route to membrane inhomogeneity.

To dissect the contribution of PEG-lipid crowding to this inhomogeneity, we constructed a phase diagram using SLBs containing varying mole percentages of PEG5000-PE (Figure 4, top row). We also compared PEG headgroups of different molecular sizes (Figures 4 and S5). From these measurements, the inhomogeneity visualized by nanodisc-delivered Texas Red-DHPE was found to depend directly on both the fraction and the chain length of PEG-lipids. Here, the PEG surface density serves as the relevant control parameter for the transition, with higher surface densities corresponding to stronger crowding and enhanced excluded-volume interactions. Notably, smaller PEG chains required higher molar fractions to induce membrane inhomogeneity, whereas longer chains produced the same effect at lower concentrations. This reciprocal relationship defines a surface-density–controlled condensation threshold of the membrane-anchored PEG layer.

FRAP analysis of SLBs containing 0.75 mol% PEG5000-PE revealed slow and spatially heterogeneous fluorescence recovery, consistent with restricted lipid exchange between mobile and immobile regions. The immobile regions were labeled equivalently with multiple fluorophores (Figure 3B), confirming that the observed behavior is not dye-specific. Control experiments further demonstrated that this crowding-induced behavior is not driven by PE lipids themselves, as SLBs with comparable PE content but lacking PEG groups remained homogeneous (Figures 4 and S6). Together, these data indicate that increasing the surface density of PEG-lipids drives a condensation transition of the PEG layer, which subsequently induces lateral membrane inhomogeneity.

We propose that this PEG-driven condensation above the SLB couples to the underlying bilayer via lipid anchorage, thereby inducing spatially heterogeneous lipid organization and diffusion, rather than intrinsic lipid–lipid phase separation, analogous to protein-driven membrane reorganization reported previously (78, 79). While crowding-induced phase transitions are well documented in solution (14–16), our work represents the first systematic demonstration of an analogous phenomenon occurring in two dimensions, in which a phase transition of membrane-anchored macromolecules reorganizes the lipid bilayer without inducing lipid demixing. The schematic in Figure 3 summarizes this mechanism, in which polymer crowding above the SLB drives membrane phase separation through tethered coupling.

Given the widespread use of PEGylated lipids in diverse membrane systems and in lipid nanoparticle (LNP)–based RNA vaccines, their intrinsic crowding effects on membrane organization warrant careful consideration. Here, nanodelivery serves as an effective strategy for introducing defined amounts of lipids into supported lipid bilayers (SLBs), enabling the detection of otherwise marginal phase separation events. Although the observed membrane inhomogeneity arises from condensation of membrane-anchored PEG groups rather than from intrinsic lipid–lipid phase separation, it nonetheless exerts a direct and measurable influence on membrane dynamics.

### Crowding-Driven Compartmentalization and Restricted Diffusion in Membranes

At moderate PEG5000-PE densities, the immobile domains form barrier-like structures that render the SLB reminiscent of the picket-fence model of the plasma membrane. The stepwise fluorescence recovery observed in FRAP experiments mirrors the spatial distribution of these immobile domains (Figures 5A, 5B and S7), indicating that two-dimensional diffusion is restricted within confined regions delineated by the barriers. Single-molecule tracking further reveals that fluorescently labeled, membrane-anchored proteins diffuse within compartments bounded by the immobile phase (Figure S8), paralleling the partitioning of the plasma membrane by cytoskeleton-associated proteins in the canonical picket-fence model.

At higher PEG-lipid fractions, condensation of the PEG layer covers most of the membrane surface, and the dense network of tethered lipids collectively yields a largely immobile bilayer (Figures 1A, 4, and 5C). Together, these findings demonstrate that molecular crowding alone can drive lateral inhomogeneity in membranes, producing organization reminiscent of the picket-fence architecture described for living cells.

### Molecular Crowding as a General Principle Governing Membrane Organization and Signaling

To further elucidate the generality of membrane-associated crowding on phase transitions, we induced the condensation of fluorescent proteins by increasing their surface density on the membrane (see Figure 6). In this case, protein surface density plays the same physical role as PEG surface density, acting as a crowding parameter that drives condensation once a critical threshold is exceeded. Unlike prior studies in which phase separation was initiated by soluble crowding agents, membrane recruitment confines proteins that freely diffuse in three dimensions in solution to lateral diffusion within the two-dimensional membrane plane, thereby amplifying effective concentration and excluded-volume interactions (70, 80). As surface density increases, this enhanced crowding drives a condensation transition among membrane-anchored proteins, which in turn produces spatial inhomogeneity in the membrane environment.

Notably, in this protein-based system, the macromolecules are anchored exclusively to the outer surface of the SLB via click chemistry and are physically separated from the solid support by the lipid bilayer and hydration layer. Nevertheless, FRAP measurements reveal similarly slow and spatially heterogeneous recovery, paralleling the behavior observed for PEG-crowded membranes. This correspondence demonstrates that crowding and condensation of membrane-anchored macromolecules alone are sufficient to generate reduced mobility and apparent immobility, independent of direct substrate coupling.

Furthermore, similar to PEG-lipid–driven transitions, the protein condensates were effectively labeled via nanodisc-mediated lipid delivery (Figure 7). The correspondence between PEG-lipid– and protein-driven phase transitions suggests a shared physical origin, in which molecular crowding, arising from PEG polymers or membrane-anchored proteins, governs lateral organization on membranes. These results also highlight nanodelivery as a powerful approach for visualizing otherwise hidden mesoscale heterogeneity on membrane surfaces.

Taken together, the PEG- and protein-crowding systems demonstrate that slow fluorescence recovery and immobile regions primarily reflect crowding-induced condensation of macromolecules at the membrane interface. While substrate coupling may influence absolute diffusion rates in glass-supported bilayers, it is not required to generate the qualitative features of immobility and compartmentalization observed here.

Many membrane-associated signaling pathways involve protein phase transitions that are coupled to the recruitment of downstream effectors to the membrane, yet our findings reveal that such transitions can also arise independently through local crowding effects. The multivalent interactions commonly observed in biological systems, which are often linked to protein phase transitions, serve as critical regulators that initiate and organize molecular assemblies at the membrane interface. As proteins are recruited to the membrane, the reduction of translational dimensionality from three to two dimensions, together with reduced diffusivity of membrane-tethered molecules, elevates local protein concentration, thereby amplifying crowding and promoting cooperative phase behavior among signaling proteins. Given that macromolecular crowding is an intrinsic feature of cellular membranes, our results underscore its general importance in modulating membrane-associated phase transitions and signaling processes.

Many membrane-associated signaling pathways involve protein phase transitions that are coupled to the recruitment of downstream effectors to the membrane, yet our findings reveal that such transitions can also arise primarily through crowding associated with high surface density, without requiring specific multivalent interactions. The reduction of translational dimensionality from three to two dimensions, together with reduced diffusivity of membrane-tethered molecules, elevates local protein concentration, thereby amplifying crowding and promoting cooperative phase behavior among signaling proteins. In this sense, increasing membrane surface density functions analogously to increasing pressure in a confined system, driving condensation and emergent membrane inhomogeneity. Given that macromolecular crowding is an intrinsic feature of cellular membranes, our results establish surface-density–driven condensation as a general physical mechanism by which the crowding effect induced by membrane recruitment gives rise to membrane organization and signaling outcomes.

## Conclusion

In summary, our study systematically elucidates how crowding by membrane-tethered macromolecules dictates molecular dynamics and organization within SLBs. By tuning the density and chain length of PEG-conjugated lipids, we revealed that the PEG layer anchored above the membrane undergoes a crowding-driven condensation transition, which in turn gives rise to mesoscale membrane heterogeneity. At sufficiently high surface density, this transition can be interpreted in terms of a percolation-like process within the PEG layer, in which overlapping PEG coronas progressively connect to form a laterally continuous polymer network at the membrane interface. Rather than invoking a lipid phase transition, the observed membrane heterogeneity emerges as a secondary consequence of connectivity and condensation within the membrane-anchored PEG layer. Using nanodisc-mediated lipid delivery, we further demonstrated controlled incorporation of fluorescent lipids into preformed SLBs, enabling direct visualization of immobile, network-like regions associated with the percolated PEG layer. These structures reflect spatially heterogeneous lipid mobility induced by polymer condensation and connectivity at the membrane interface and recapitulate key features reminiscent of the picket-fence model. Moreover, analogous condensation behavior observed for membrane-anchored proteins under high surface density underscores that surface-density–driven crowding can give rise to percolation-like connectivity among membrane-tethered macromolecules, independent of specific chemical interactions. Given that macromolecular crowding is ubiquitous at cellular membranes and that PEG-functionalized lipids are widely employed in nanotechnology and pharmaceutical formulations, our findings reveal a general physical principle in which surface-confined macromolecules undergo crowding-induced condensation and connectivity transitions in which isolated clusters merge into laterally continuous networks, reorganizing membrane structure and dynamics through percolation-like network formation at membrane interfaces.

## Materials and Methods

### Materials

1,2-dioleoyl-sn-glycero-3-phosphocholine (DOPC), 1,2-dioleoyl-sn-glycero-3-phospho-L-serine (DOPS), 1,2-dioleoyl-sn-glycero-3-phosphoethanolamine-N-[methoxy(polyethylene glycol)-5000] (ammonium salt) (PEG5000-PE), 1,2-dioleoyl-sn-glycero-3-phosphoethanolamine-N-[methoxy(polyethylene glycol)-3000] (ammonium salt) (PEG3000-PE), 1,2-dioleoyl-sn-glycero-3-phosphoethanolamine-N-[methoxy(polyethylene glycol)-2000] (ammonium salt) (PEG2000-PE), 1,2-dioleoyl-sn-glycero-3-phosphoethanolamine-N-[methoxy(polyethylene glycol)-1000] (ammonium salt) (PEG1000-PE), and 1,2-dioleoyl-sn-glycero-3-phosphoethanolamine-N-[methoxy(polyethylene glycol)-350] (ammonium salt) (PEG350-PE) were purchased from Avanti Polar Lipids (Alabaster, AL). 1,2-dioleoyl-sn-glycero-3-[(N-(5-amino-1-carboxypentyl)iminodiacetic acid)succinyl] (Ni²[-NTA-DOGS; nickel salt) was also obtained from Avanti Polar Lipids. Texas Red-DHPE and fluorescein-DHPE (FITC-DHPE) were purchased from Thermo Fisher Scientific. Sulfuric acid (H[SO[) and hydrogen peroxide (H[O[) were purchased from Honeywell Fluka. Tris-buffered saline (TBS) was purchased from Protech Technology. Alexa Fluor 647 NHS ester was purchased from Lumiprobe, and bovine serum albumin (BSA) was obtained from Sigma-Aldrich.

## Preparation of small unilamellar vesicles (SUVs)

Small unilamellar vesicles (SUVs) were prepared from lipid mixtures containing DOPC, PEG-phospholipids of different chain lengths (0.75–3 mol%), FITC-DHPE (0.5 mol%), and DOPS (2 mol%) dissolved in chloroform. In the single-molecule experiment using Alexa Fluor 647–labeled TEV protease, SUVs were prepared from lipid mixtures containing DOPC, PEG-phospholipids (1 mol%), FITC-DHPE (0.5 mol%), and Ni²[-NTA-DOGS (4 mol%) dissolved in chloroform. The lipid mixtures were evaporated using a rotary evaporator at 40 °C for 3 min to remove chloroform and form a dry lipid film, which was then further dried under a gentle stream of N[ for 8 min. The dried lipid film was rehydrated with 2 mL of deionized water, followed by vortexing and gentle pipetting to resuspend the lipids and form multilamellar vesicles of varying sizes. SUVs were subsequently obtained by tip sonication to disperse the vesicles formed in the previous step.

## Preparation of nanodiscs

### Expression and purification of MSP

Membrane scaffold protein (MSP1D1, referred to as MSP for simplicity) was expressed and purified with minor modifications to a previously described protocol (55). Briefly, the pET-28a plasmid containing the MSP gene (Addgene) was freshly transformed into *E. coli* BL21(DE3) cells (Agilent) and grown on kanamycin-supplemented agar plates (30 µg/mL) at 37 °C. A single colony was then transferred into 10 mL of Terrific Broth (TB) containing 30 µg/mL kanamycin and grown until the optical density at 600 nm (OD600) reached 0.6–0.8. The culture was scaled up by inoculating 0.6 L of TB with 30 µg/mL kanamycin, grown at 37 °C, and induced with 1 mM IPTG for 4 hours at 28 °C when OD600 reached approximately 2.0–2.5. The cells were harvested by centrifugation and stored at −80 °C.

For purification, the cell pellets were resuspended in 30 mL of buffer containing 20 mM sodium phosphate, 0.1 M NaCl, 1% Triton X-100, 10 mM MgSO_4_, pH 7.4, supplemented with 10 µg/mL DNase I and 300 µL of 0.1 M phenylmethylsulfonyl fluoride (PMSF) in ethanol. The suspension was sonicated, and cell debris was removed by centrifugation at 12,800 × g for 50 minutes. The supernatant was loaded onto a 5 mL HisTrap HP column (GE Healthcare) pre-equilibrated with 20 mM sodium phosphate, 0.1 M NaCl, and 1% Triton X-100, pH 7.4. The column was sequentially washed with the following buffers: (i) 40 mM Tris-HCl, 0.3 M NaCl, 1% Triton X-100, pH 8.0 (25 mL); (ii) 40 mM Tris-HCl, 0.3 M NaCl, 50 mM sodium cholate (SC), pH 8.0 (25 mL); (iii) 40 mM Tris-HCl, 0.3 M NaCl, 40 mM imidazole, pH 8.0 (25 mL). MSP was eluted with 40 mM Tris-HCl, 0.3 M NaCl, and 0.4 M imidazole, pH 8.0. The eluted protein was buffer-exchanged into MSP buffer (20 mM Tris-HCl and 0.1 M NaCl, pH 7.4) and concentrated to ∼10 mg/mL using a 10 kDa MWCO centrifugal concentrator (Amicon). The concentration of MSP was determined by measuring absorbance at 280 nm, using an extinction coefficient of 21,430 M^-1^ cm^-1^.

### Preparation of nanodisc samples

Nanodiscs were prepared following an established protocol with defined reconstitution ratios: lipid/MSP at 65:1 and sodium cholate (SC)/lipid at 2:1. For lipid film preparation, 4% fluorophore-conjugated lipids (FITC-DHPE or Texas Red-DHPE; Thermo Fisher) were mixed with pure DOPC (Avanti) in chloroform. This composition ensures that over 99.5% of the nanodiscs contain at least one fluorescent lipid in the bilayer, as calculated from a binomial distribution. The lipid mixtures were dried under a gentle nitrogen flow for 30 minutes and solubilized by adding SC, followed by sonication. MSP was then added to the solubilized lipid solution, and the mixture was incubated on ice for 30 minutes.

To assemble nanodiscs, SM-2 Biobeads (1 g/mL; Bio-Rad) were added to the mixture and incubated at 4 °C for 5 hours. Biobeads were removed by centrifugation, and the resulting fractions were purified using a Superdex 200 10/300 GL gel filtration column (GE Healthcare). Purified nanodiscs were buffer-exchanged into nanodisc buffer (50 mM HEPES and 0.1 M NaCl, pH 7.0) and concentrated using Amicon Ultra-50K centrifugal filter units. Nanodisc concentrations were determined by absorbance at 280 nm.

## TEV purification

The pRK508 plasmid containing the TEV S219V mutant fused to an N-terminal His[ tag and an MBP fusion tag with a tobacco etch virus (TEV) cleavage site was transformed into E. coli BL21 (DE3). The successfully transformed colonies were inoculated into 1 L of Terrific Broth (TB) medium and incubated at 37 °C until the optical density at 600 nm (OD[[[) reached 0.6. Protein expression was induced with 1 mM IPTG, and the culture was further incubated at 37 °C for 3 h. The culture was centrifuged at 6000 × g for 20 min, and the resulting cell pellet was resuspended in 50 mL of buffer A (50 mM HEPES, 1 M NaCl, 20 mM imidazole, 10% glycerol, pH 7.5).

The resuspended cells were lysed by sonication, followed by centrifugation at 15 000 × g for 30 min to remove cell debris. The supernatant was loaded onto a HisTrap FF column (GE Healthcare) pre-equilibrated with buffer A. After removing non-target proteins, TEV protease was eluted with buffer C (50 mM HEPES, 50 mM NaCl, 400 mM imidazole, 10% glycerol, pH 7.5). The eluted fractions were concentrated to 5 mL and buffer-exchanged into buffer B (50 mM HEPES, 50 mM NaCl, 20 mM imidazole, 10% glycerol, pH 7.5) using an Amicon Ultra centrifugal filter unit (10 kDa molecular weight cutoff [MWCO]; Millipore). The concentrated sample was applied to a HiTrap SP FF column (GE Healthcare) to further purify TEV protease by adjusting the NaCl concentration through a gradient between buffer B and buffer D. The fraction containing TEV was eluted at approximately 25% buffer D (50 mM HEPES, 400 mM NaCl, 20 mM imidazole, 10% glycerol, pH 7.5). The eluted protein was concentrated and buffer-exchanged into the size-exclusion buffer using an Amicon Ultra centrifugal filter unit (10 kDa MWCO). The concentrated sample was subsequently loaded onto a Superdex 75 10/300 GL column (GE Healthcare) equilibrated with the size-exclusion buffer (0.1 M PB, pH 8.0) . Fractions containing TEV protease were collected, and the protein concentration was determined by UV–Vis spectroscopy.

The purified TEV protease was quick-frozen in liquid nitrogen and stored at −80 °C for subsequent labeling reactions.

## eGFP purification

The enhanced green fluorescent protein (eGFP) sequence with an N-terminal His[ tag and a TEV protease cleavage site was cloned into the pET-28 plasmid. The plasmid was transformed into E. coli BL21 (DE3) using the heat-shock method. Transformed colonies were inoculated into 1 L of Terrific Broth (TB) medium and cultured at 37 °C until the optical density at 600 nm (OD[[[) reached ∼0.6. Protein expression was induced with 1 mM isopropyl β-D-1-thiogalactopyranoside (IPTG), and the culture was further incubated at 37 °C for 2 h. The cells were harvested by centrifugation at 6000 × g for 20 min, and the pellet was collected for purification.

The cell pellet was resuspended in 50 mL of Ni–NTA buffer (20 mM Tris–HCl, 500 mM NaCl, 20 mM imidazole, 10% glycerol, pH 8.0) by vortexing and lysed by sonication. The cell debris was removed by centrifugation at 15 000 × g for 30 min. The supernatant was loaded onto a HisTrap FF column (GE Healthcare) pre-equilibrated with Ni–NTA buffer. After washing to remove non-target proteins, eGFP was eluted using Ni–elution buffer (20 mM Tris–HCl, 500 mM NaCl, 500 mM imidazole, 10% glycerol, pH 8.0). The eluted protein was concentrated to 500 µL and buffer-exchanged into size-exclusion buffer (0.1 M phosphate buffer, pH 8.0) using an Amicon Ultra centrifugal filter unit (10 kDa MWCO; Millipore). The concentrated sample was then loaded onto a Superdex 75 10/300 GL column (GE Healthcare) equilibrated with the same buffer. Fractions containing eGFP were collected, and protein purity was verified by SDS–PAGE. After determining the concentration, the purified eGFP was aliquoted, flash-frozen in liquid nitrogen, and stored at −80 °C.

## mNeonGreen purification

The pET-28 vector containing the mNeonGreen sequence with an N-terminal His[ tag and a TEV protease cleavage site was used for protein expression. The expression and purification procedures were identical to those described for eGFP.

## Preparation of Cys-TEVcut-mScarlet

The DNA construct His[-Factor Xa cut site-Cys-TEVcut-mScarlet was generated by gene cloning. The His[-tag facilitated purification, the Factor Xa site enabled tag removal, the cysteine residue provided a reactive thiol for coupling with MCC lipids, and mScarlet served as the fluorescent probe (Avanti Polar Lipids, cat. 780201P). The construct in pET28 was transformed into BL21 (DE3) cells, grown in 1 L of Terrific Broth at 37 °C, and induced with 1 mM IPTG for 3 h once OD[[[ reached 0.6–0.8. Cells were harvested (6000 × g, 20 min, 4 °C), lysed by sonication in 20 mM Tris–HCl buffer (500 mM NaCl, 20 mM imidazole, 10 % glycerol, pH 8.0), and clarified (15 000 × g, 30 min, 4 °C). The protein was purified using HisTrap FF and Superdex 75 10/300 columns (GE Healthcare). The His[ tag was removed with Factor Xa protease for 18 h at 4 °C, and the cleaved sample was reapplied to the HisTrap column to remove uncleaved protein and free tag. The flow-through was collected, aliquoted, flash-frozen, and stored at −80 °C for subsequent imaging experiments.

## Experiments using click chemistry

For related experiments, SLBs were prepared using SUVs composed of 93 mol% DOPC, 5 mol% DBCO-PE, and 2 mol% DOPA on piranha-etched glass substrates. DBCO-PE lipids were conjugated with mScarlet through copper-free click chemistry. SUVs were prepared by mixing lipids at a molar ratio of DOPC:DBCO-PE:DOPA = 93:5:2 in chloroform. The mixture was evaporated using a rotary evaporator for 3 min at 40 °C and dried under N[ for 15 min to form a lipid film, which was resuspended in 2 mL deionized water and sonicated to yield uniform SUVs.

Glass substrates were cleaned with piranha solution (H[SO[:H[O[ = 3:1 v/v), rinsed thoroughly, dried with N[ (g), and assembled into flow chambers. SUVs were mixed with TBS buffer (1:1 v/v) and incubated for 20 min to form SLBs. Cys-mScarlet (25–30 µM) was reacted with eight equivalents of Azide-PEG[-NHS ester in PBS at 37 °C for 4 h to introduce azido groups. The modified mScarlet was then incubated with DBCO-PE SLBs at room temperature for 10 h to achieve copper-free click conjugation. By varying the protein concentration, the surface density of mScarlet anchors on the SLBs was controlled. Unbound proteins were removed by washing with PBS buffer prior to imaging. Fluorescence images were acquired using a Nikon ECLIPSE Ti2-E microscope equipped with a 100× 1.49 NA oil-immersion TIRF objective.

## Experiments using maleimide chemistry

SLBs were prepared using SUVs composed of 94 mol% DOPC, 4 mol% Ni²[-NTA DOGS, and 2 mol% PE-MCC on piranha-etched glass substrates. PE-MCC lipids were labeled with the fluorescent protein mScarlet through maleimide chemistry. SUVs were prepared by mixing lipids at a molar ratio of DOPC:Ni²[-NTA DOGS:PE-MCC = 94:4:2 in chloroform. The lipid mixture was evaporated using a rotary evaporator for 3 min at 40 °C to remove chloroform and form a dried lipid film, followed by further drying under N[ for 15 min. The lipid film was resuspended in 2 mL of deionized water by vortexing and pipetting. A uniform SUV suspension was obtained by tip sonication to disrupt larger vesicles.

Glass substrates were treated with piranha solution (H SO :H O = 3:1 v/v), thoroughly rinsed with deionized water, blow-dried with N[ (g), and assembled into flow chambers. SUVs were mixed with TBS buffer (1:1 v/v) and injected into the chamber for 20 min to form SLBs. Residual defects were blocked by incubation with 1 mg mL[¹ BSA for 15 min. Cys-mScarlet (250 nM) was then incubated with the SLBs for 20 min to react with PE-MCC lipids via maleimide coupling. Unbound proteins were removed by washing with TBS buffer. Fluorescence imaging was performed using a Nikon ECLIPSE Ti2-E microscope equipped with a 100× 1.49 NA oil-immersion TIRF objective. Uniform mScarlet fluorescence was observed from SLBs containing 94 mol% DOPC, 4 mol% Ni²[-NTA DOGS, and 2 mol% PE-MCC, with no phase separation between PC and PE lipids (Figure S6A).

The mobility and recovery of the bilayer were characterized by fluorescence recovery after photobleaching (FRAP). After exposure to laser irradiation at high power for 1 min within a small region, fluorescent images were collected every 10 s to monitor recovery (Figure S6B). The bleached area showed rapid and uniform recovery from the edges toward the center without a grid-like pattern, indicating free diffusion of both PC and PE lipids across the SLBs.

## Protein labeling

Protein fluorescence labeling was performed at a 1:1 molar ratio of dye to protein. Alexa Fluor 647 NHS ester was dissolved in anhydrous DMSO to prepare a 2 mg/mL dye stock solution. TEV was concentrated such that the protein-to-dye volume ratio was maintained at 9:1. After mixing the protein and dye solutions, the labeling reaction was carried out at room temperature for 1 h, followed by incubation at 4 °C for 8 h. The precipitate was removed from the labeled sample by centrifugation at 15 000 × *g*, and the supernatant containing labeled TEV was collected. Unreacted dye was removed using an Amicon Ultra centrifugal filter unit (10 kDa MWCO; Millipore), and the sample was subsequently concentrated. The concentrated sample was further purified by size-exclusion chromatography using a Superdex 75 (GE Healthcare) column. The protein concentration and labeling efficiency were determined by UV–Vis spectroscopy. Finally, aliquots of the labeled protein were quick-frozen in liquid nitrogen and stored at −80 °C for future use.

## Supported lipid bilayers (SLBs)

Supported lipid bilayers (SLBs) were prepared by vesicle fusion on a glass substrate. Glass coverslips (thickness 170 ± 5 µm; Ibidi) were etched with piranha solution (a 3:1 v/v mixture of H[SO[ and 30% H[O[) for 8 min. The etched glass was then assembled into a flow chamber (Sticky-Slide VI 0.4; Ibidi). A 1:1 volume mixture of the SUV suspension and TBS buffer was prepared and injected into the chamber, followed by a 20 min incubation to allow SLB formation. After incubation, excess vesicles were removed by washing with 900 µL of TBS buffer. Next, 0.5 µL of nanodisc solution was mixed with 99.5 µL of TBS, added to the prepared SLB, and incubated for 5 min. Excess nanodiscs were then washed off with 900 µL of TBS buffer.

In the single-molecule TEV experiment, a 1:1 mixture of the SUV suspension and TBS buffer was injected into the chamber and incubated for 20 min to form SLBs. Afterward, excess vesicles were removed by washing with 900 µL of TBS buffer. The SLBs were then blocked with 1 mg/mL BSA for 15 min, followed by washing with 900 µL of TBS to remove unbound BSA. Subsequently, 0.5 µL of nanodisc solution mixed with 99.5 µL of TBS was added to the SLB, incubated for 5 min, and washed out with 900 µL of TBS buffer. Finally, the SLB was incubated with TEV protease (0.02 µM) for 5 min, followed by a final wash with 900 µL of TBS buffer. At this stage, the His-tag on TEV allowed its recruitment to the nickel nitrilotriacetic acid (Ni²[–NTA) groups on the SLB.

## TIRF Microscopy

Imaging was carried out using a Nikon Eclipse Ti2 inverted microscope equipped with a total internal reflection fluorescence (TIRF) system and an electron-multiplying CCD (EMCCD) camera (iXon Life 897; Andor Technology). The TIRF microscopy setup utilized a Nikon 100×/1.49 NA oil-immersion objective, along with a TIRF illuminator, Perfect Focus System, motorized stage, and U-N4S four-laser unit (Nikon) as the excitation source. This configuration included solid-state lasers with wavelengths of 488 nm, 561 nm, and 640 nm, controlled by an integrated acousto-optic tunable filter (AOTF). The laser power was adjusted to 5.2 mW (488 nm), 6.9 mW (561 nm), and 7.8 mW (640 nm), as measured before the objective with the field aperture fully open. Fluorescence filtering was performed using a 405/488/561/638 nm Quad TIRF filter set (Chroma Technology Corp.).

## Imaging Analysis

Using TrackMate (ImageJ/Fiji plugin), particles were detected using the Laplacian of Gaussian (LoG) detector with a threshold of 40 and an estimated blob diameter of 1.2 µm. The detected spots were then linked into trajectories using the Simple LAP (Linear Assignment Problem) tracker, with a maximum linking distance of 1.2 µm and allowing for up to four frames of gap closing to account for temporary signal loss.

## Author Contributions

R.-H.B., H.-Y.T., Y.-W.C. and C.-W.L. designed research; R.-H.B., H.-Y.T., C.-C.C., H.-J.T., C.-K.L., C.-J.K., and W.-T.W. performed research; C.-M.J. enhanced the sensitivity of the optical systems; R.-H.B., H.-Y.T., Y.-W.C. and C.-W.L. analyzed and discussed data; and R.-H.B., H.-Y.T., Y.-W.C. and C.-W.L. wrote the paper.

