## Supplementary material for "Nanodisc-Mediated Visualization of Crowding-Induced Condensation and Membrane Reorganization in Two-Dimensional Membrane Environments": SI

#### Title:

### Supporting Information

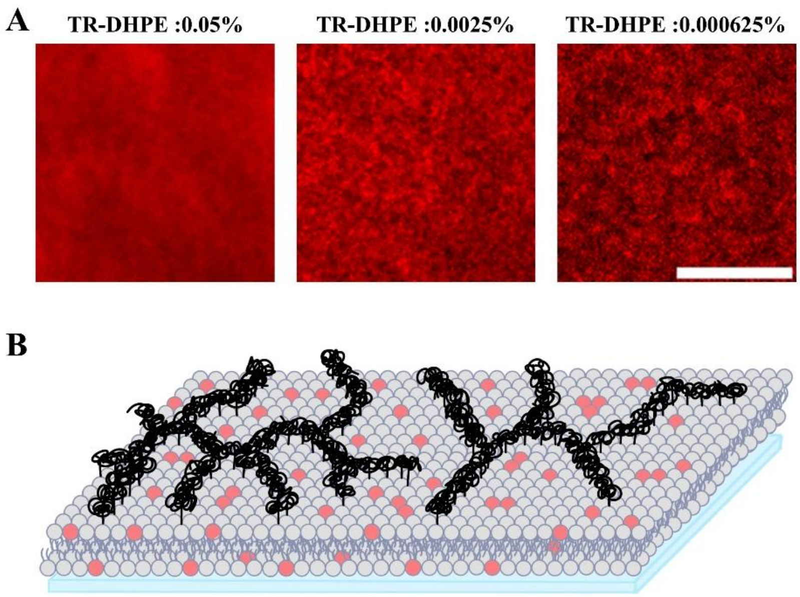


Figure S1. Fluorescence images of SLBs containing 0.75% PEG5000-PE and varying percentages of TR-DHPE. (A) The SLB appears homogeneous and remains mobile, as shown by images with TR-DHPE concentrations ranging from 0.05% to 0.006%. Inhomogeneous features cannot be clearly observed by this method, in contrast to the membrane inhomogeneity revealed by nanodelivery. (Scale bar = 8 μm) (B) The cartoon schematic illustrates that TR-DHPE lipids are uniformly distributed without preference between the mobile and immobile phases on the SLB.

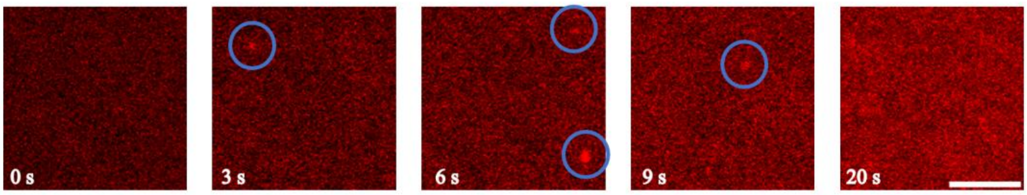


Figure S2. Fluorescent and nonfluorescent lipids are efficiently unloaded onto the SLB via nanodelivery. Nanodiscs containing 4% TR-DHPE and 96% DOPC collide with the SLB and are captured by TIRF microscopy, appearing as bright spots (blue circles) that mark lipid unloading events. The entire SLB becomes fluorescent within approximately 20 seconds. In the absence of PEG-induced crowding, the SLB exhibits homogeneous fluorescence across the field of view.

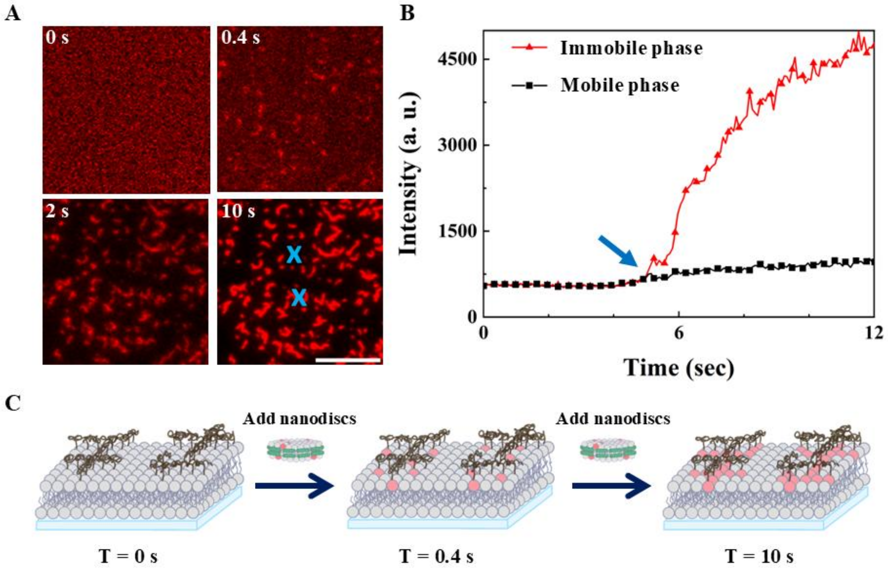


Figure S3. The immobile phase on the SLB containing 0.75% PEG5000-PE is selectively labeled by nanodelivery using nanodiscs loaded with TR-DHPE. (A) Time-lapse fluorescence snapshots show that TR-DHPE lipids are rapidly unloaded onto the immobile regions within 10 seconds after nanodisc addition. (Scale bar = 8 μm) (B) The fluorescence intensity over time at the mobile (black) and immobile (red) regions, corresponding to the blue cross marks in panel A, is plotted. The blue arrow indicates the time point of nanodisc addition (t = 0 s). (C) The schematic illustrates that Texas Red-DHPE and other nanodisc lipids preferentially unload onto the immobile PEG-lipid phase, resulting in enhanced fluorescence from these regions.

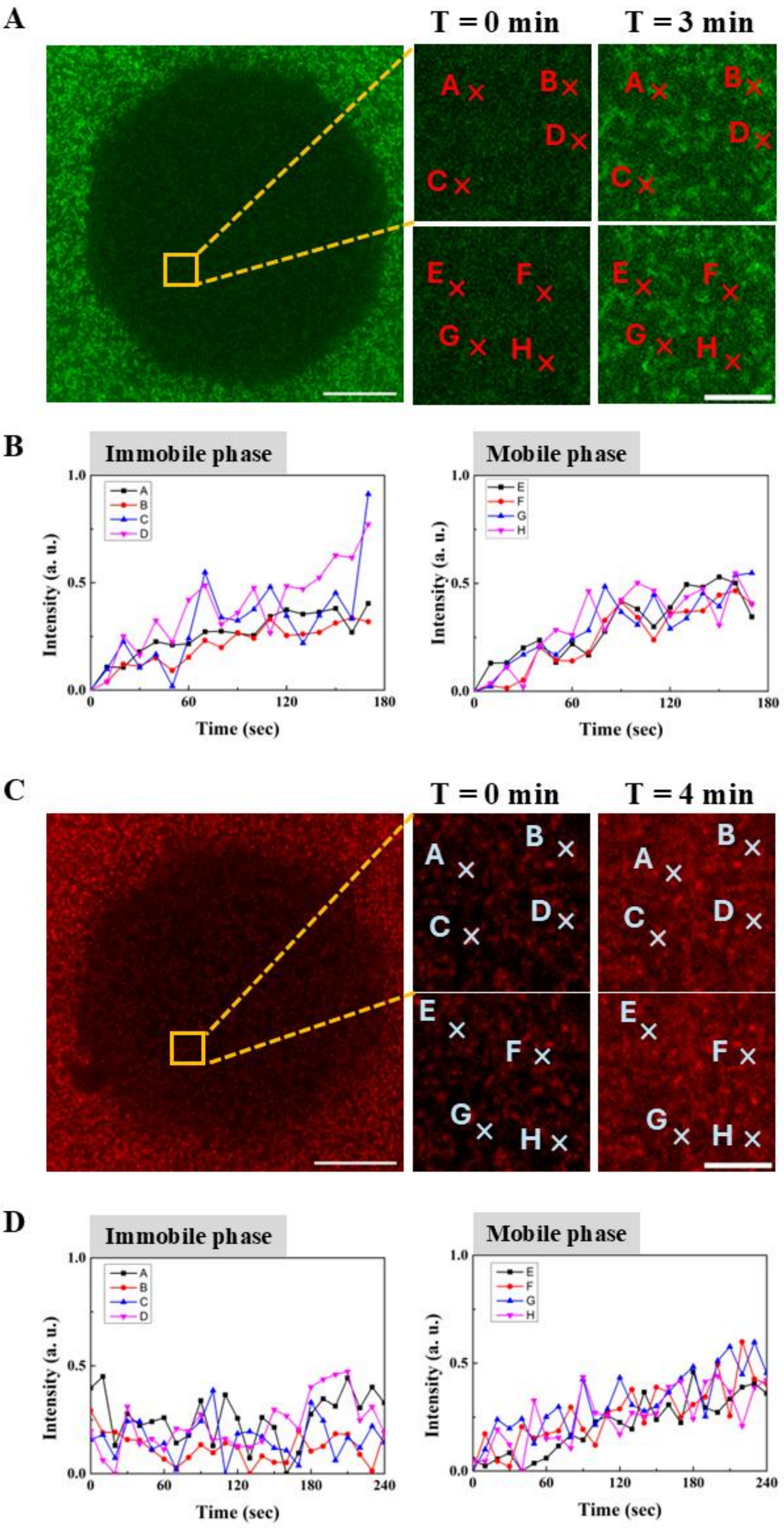


Figure S4. FRAP experiments were conducted on SLBs containing 0.75% PEG5000-PE to examine lipid exchange between the mobile and immobile phases. The crowding-induced phase transition was visualized by nanodelivery using two different fluorescent lipids, FITC-DHPE and Texas Red-DHPE. Fluorescence recovery at mobile and immobile regions was monitored simultaneously at eight locations (A–H), with the first four corresponding to immobile regions and the last four to mobile regions. (A) The phase transition highlighted by nanodiscs loaded with FITC-DHPE was analyzed by FRAP after nanodelivery. (B) The time courses of fluorescence intensity at the eight locations shown in panel A correspond to the immobile and mobile phases. The observed fluorescence recovery in the immobile phase indicates lipid exchange between the two phases. (C) The phase transition highlighted by nanodiscs loaded with Texas Red-DHPE was analyzed by FRAP after nanodelivery. (D) The corresponding fluorescence recovery curves at the eight locations shown in panel C reveal slower recovery in the immobile phase, reflecting differences arising from the fluorescent chromophore. Together, these measurements confirm lipid exchange between the mobile and immobile phases, while also demonstrating that the extent of recovery is partially dependent on the fluorescent probe, with Texas Red-DHPE showing only subtle recovery in the immobile phase.

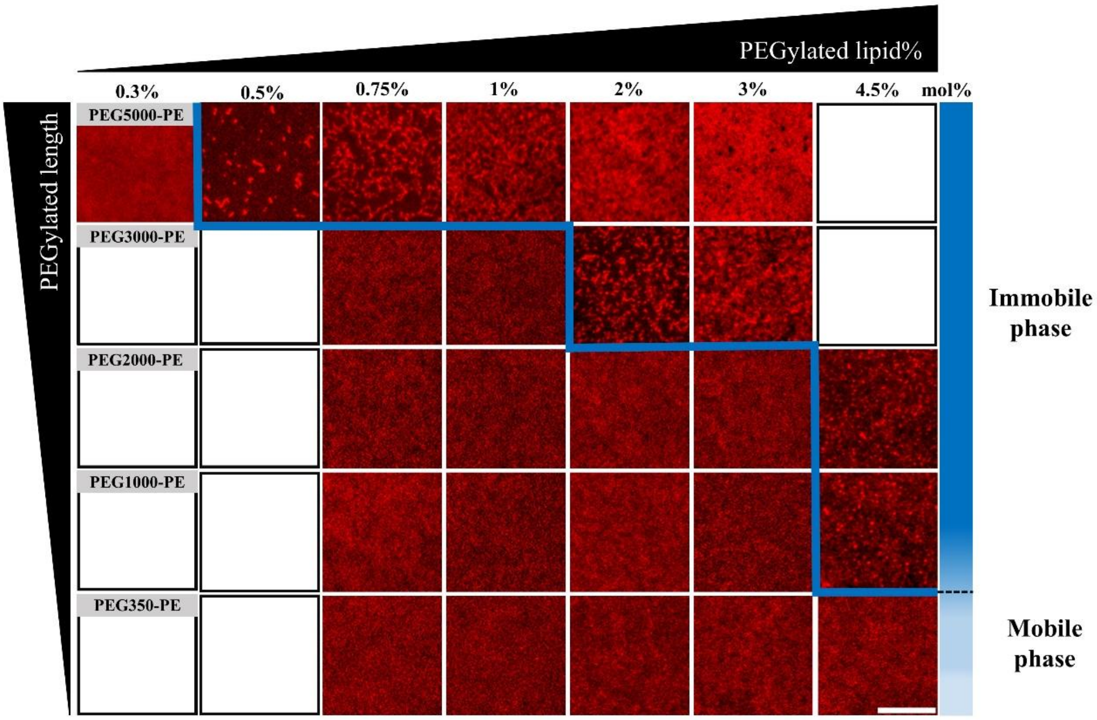


Figure S5. The more complete phase diagram illustrates that the phase transition is mediated by the molar percentage of PEG-lipid and the PEG chain length at 25 °C (scale bar = 8 μm). Once the membrane undergoes a phase transition driven by the crowding effect, the fluorescence images reveal membrane inhomogeneity, with the immobile phase preferentially labeled by fluorescent lipids delivered from nanodiscs. The area of the immobile phase increases as the PEG-lipid fraction rises, and lipids with longer PEG chains induce phase transition at lower PEG-lipid concentrations.

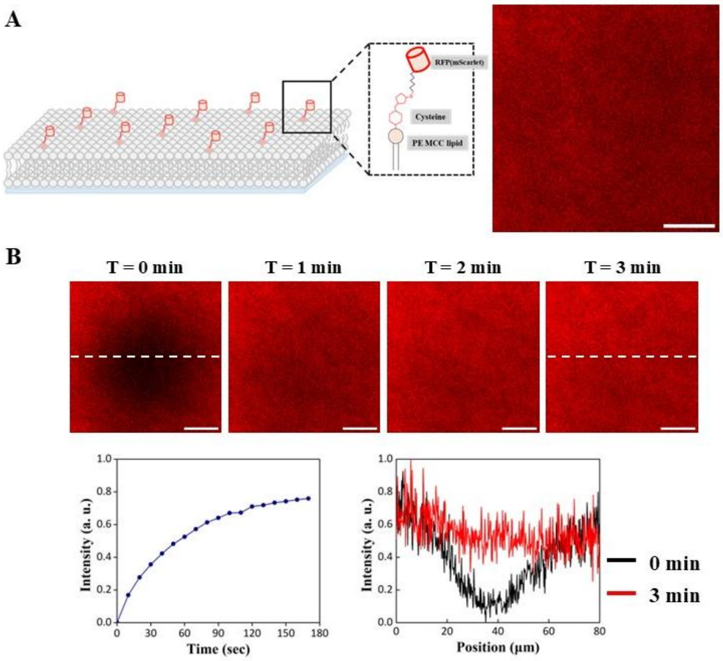


Figure S6. Uniform distribution of mScarlet across the SLB with lipid compositions similar to those used in most experiments of this study. (A) The schematic diagram on the left shows mScarlet anchored on the SLB at a surface density of approximately 450 molecules $\mu m^{-2}$. The SLB is composed of 94 mol % DOPC, 4 mol % $Ni^{2+}$–NTA DOGS, and 2 mol % PE–MCC. A cysteine residue was introduced at the N-terminus of mScarlet, allowing covalent anchoring to the SLB via maleimide coupling with PE–MCC. The fluorescence image shows a uniform distribution of mScarlet, indicating that PE–MCC lipids are homogeneously distributed across the SLB. This confirms that no phase separation arises from the PC and PE lipids alone. (B) FRAP analysis of mScarlet-labeled PE–MCC lipids in the SLB. After exposure to the laser irradiation at high power for 1 min, fluorescence images were recorded every 10 s to monitor recovery. The fluorescence recovery trace is shown on the bottom left. Fluorescence images acquired at 0, 1, 2, and 3 min are displayed at the top, and the corresponding intensity profiles along the dashed line at 0 min (black) and 3 min (red) are shown at the bottom right. The intensity of the photobleached region showed rapid and smooth recovery from the edges, confirming the free diffusion of PE lipids and indicating that PE lipids at the compositions commonly used throughout this study alone do not give rise to phase-transition-induced inhomogeneity in the SLB.

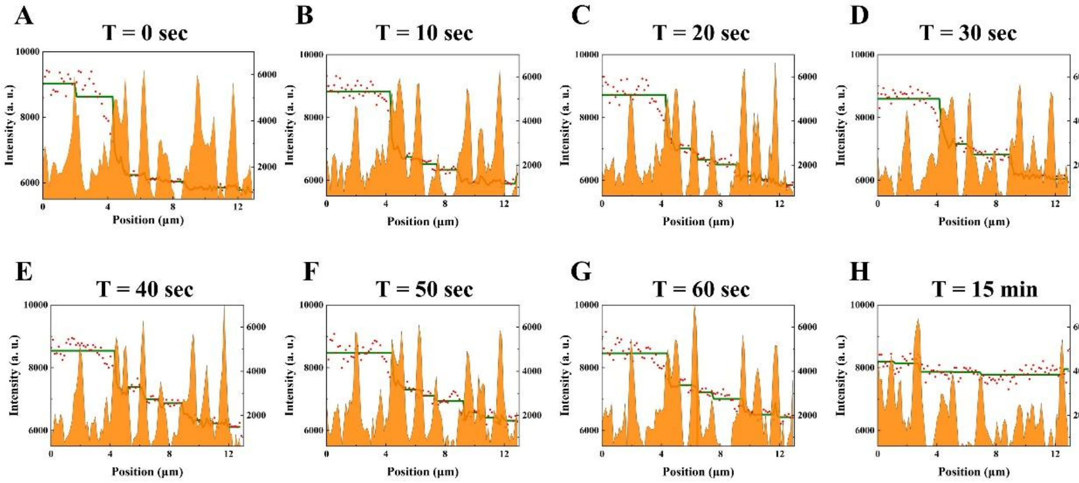


Figure S7. The immobile phase formed by PEG lipids acts as a barrier that restricts lipid diffusion on the membrane. The fluorescence intensity profiles along the dashed line in Figure 6 at different time points are plotted. The red dots represent the fluorescence intensity of FITC-DHPE, with the corresponding values indicated on the left y-axis, while the green line shows the fitted stepwise function for the red data points. The height of the orange area corresponds to the fluorescence intensity profile of the immobile phase labeled with Texas Red-DHPE along the dashed line in Figure 6, shown with reference to the right y-axis. The fluorescence intensity profiles at different time points are shown in (A) t = 0 s, (B) t = 10 s, (C) t = 20 s, (D) t = 30 s, (E) t = 40 s, (F) t = 50 s, (G) t = 60 s, and (H) t = 15 min. The fluorescence intensity profile of FITC-DHPE in the mobile phase is well fitted by a piecewise function, where the discontinuities of the green line match the orange peaks corresponding to the barriers formed by the immobile phase. The correlation between the green line and the orange peaks confirms that the inhomogeneity arising from crowding modulates membrane dynamics by limiting lipid diffusion. At longer times (t = 15 min), the relatively flat intensity profile indicates that diffusive equilibration across the barriers leads to complete fluorescence recovery after photobleaching.

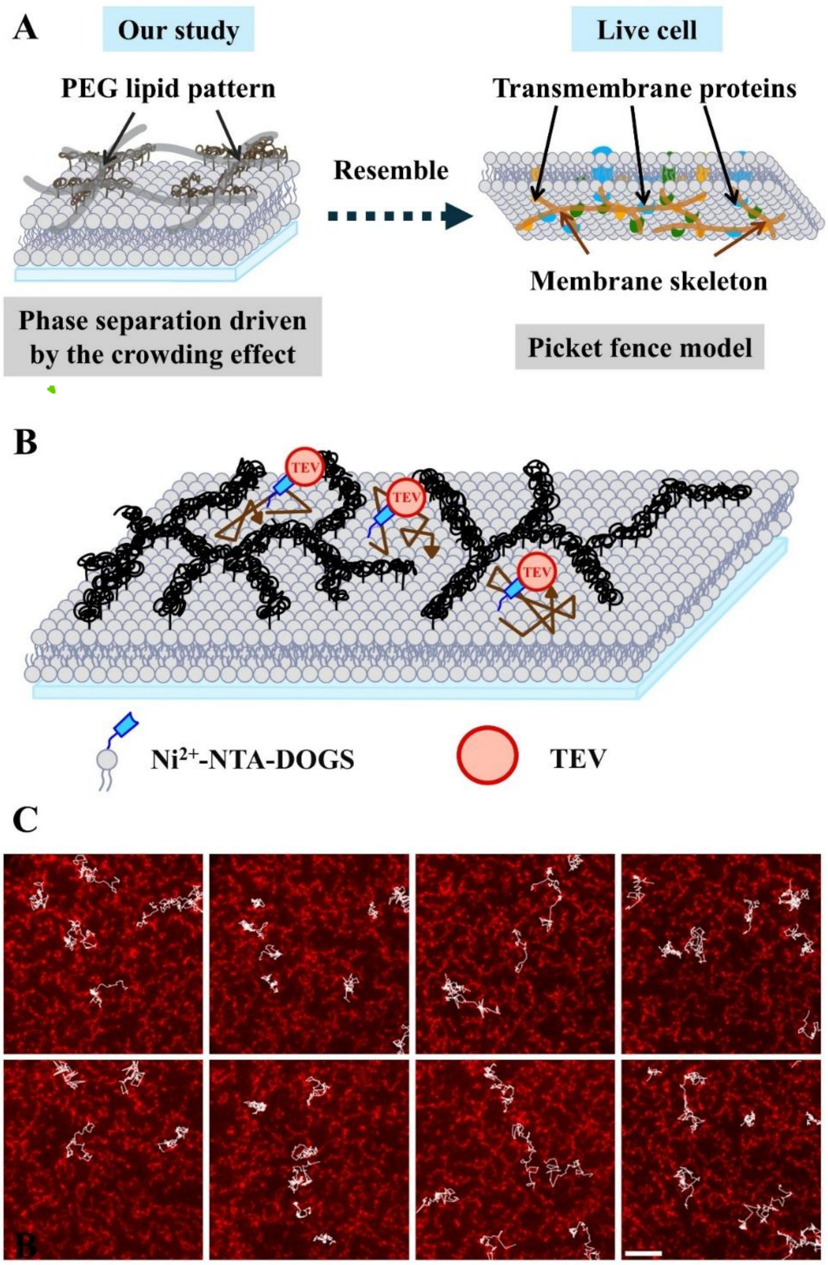


Figure S8. (A) Schematic representation of the phase transition driven by the crowding effect on the SLB that resembles the picket-fence model of the plasma membrane. In this model, the cell membrane is partitioned into microscopic domains that restrict membrane dynamics, analogous to the mobile phase confined by barriers formed by the immobile phase due to crowding in our system. (B) The fluorescence images of the immobile phase labeled with Texas Red–DHPE via nanodelivery (in red) overlaid with single-molecule tracks (in white) of TEV protein show that TEV proteins are confined within microdomains of the SLB (containing 0.75 mol% PEG5000-PE and 4 mol% Ni–NTA lipid), where phase transition is driven by the crowding effect. TEV was fluorescently labeled with Alexa Fluor 647 NHS ester (~140% labeling efficiency) and recruited to the membrane via His-tag chemistry. The fluorescence images were collected using TIRF microscopy, and the trajectories of TEV on the membrane were analyzed using the TrackMate plugin in ImageJ. The TEV trajectories are predominantly confined within the immobile phase, with only occasional events in which TEV molecules cross the barriers, resembling the picket-fence model. (Scale bar = 5 μm.)

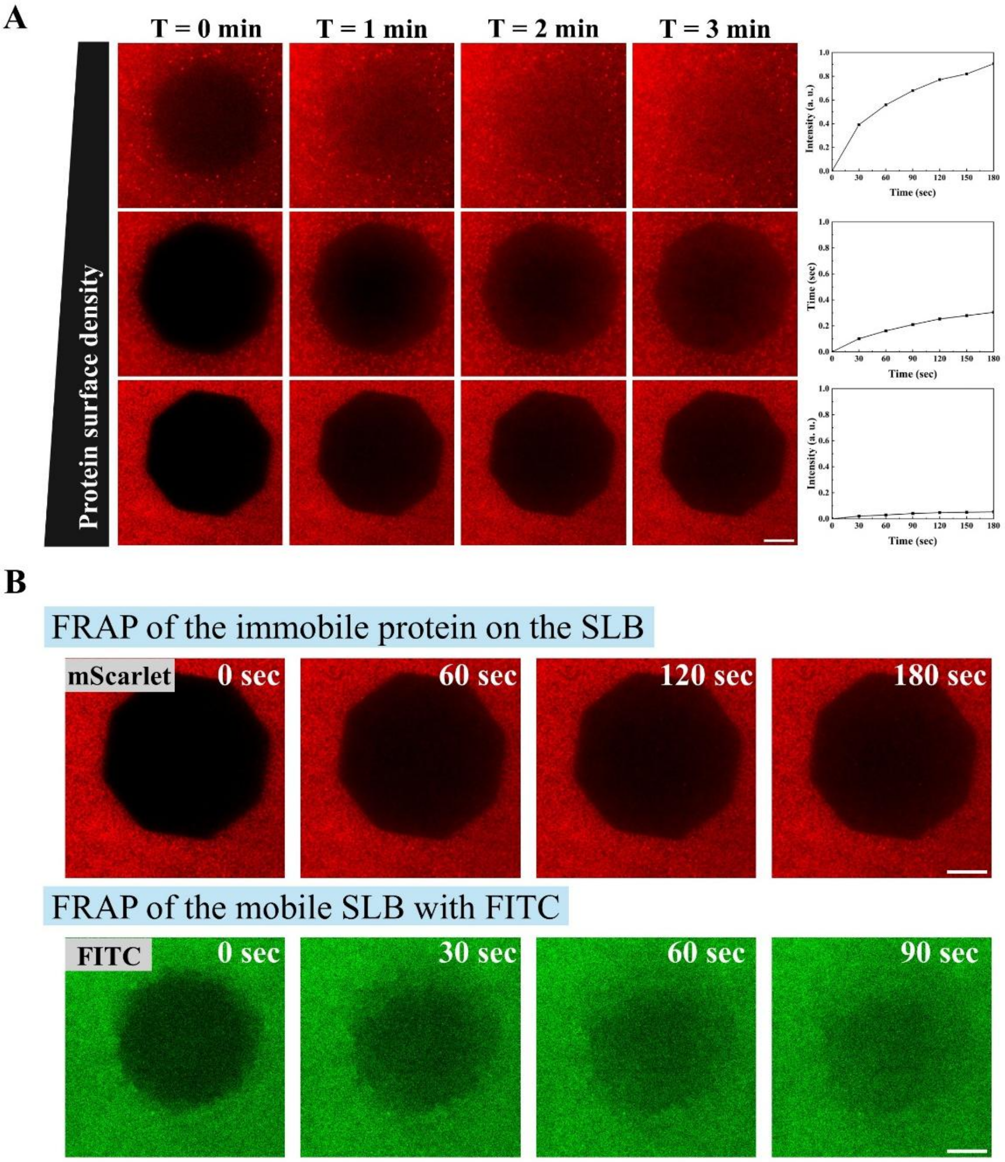


Figure S9. FRAP experiments of mScarlet anchored on SLB at different densities (from ~100 to 1800 molecules $\mu m^{-2}$) via click chemistry. (A) As the surface density of mScarlet increases, the fluorescence recovery in the FRAP experiment becomes slower, implying the formation of protein condensation. The corresponding normalized time courses of fluorescence recovery at the bleached area are shown on the right. (B) The FRAP experiments of mScarlet from the third row in part A and the corresponding FITC lipid in the same SLB are compared. The fast fluorescence recovery of FITC lipid on the membrane indicates that the membrane remains intact, while the protein forms immobile condensates on the membrane. (Scale bar = 10 μm.)

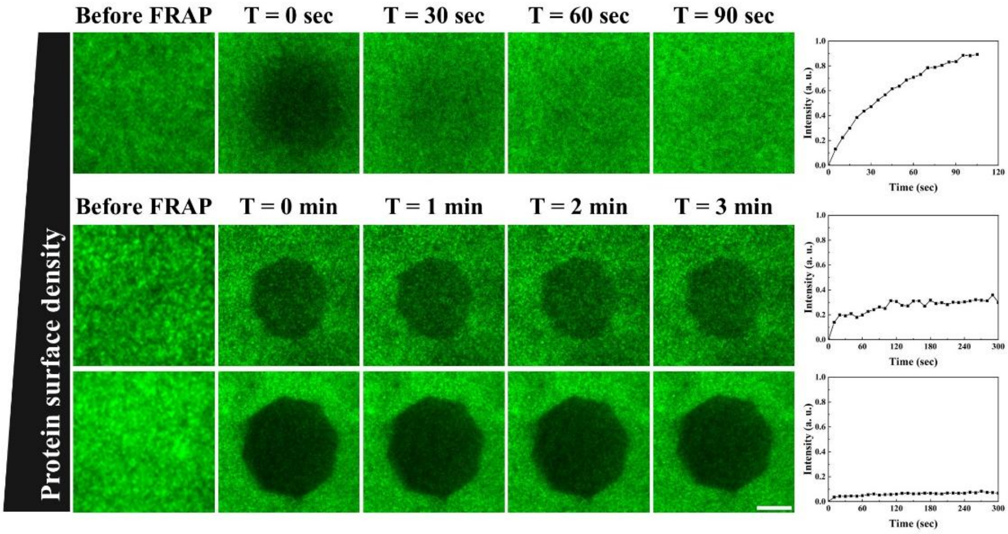


Figure S10. FRAP experiments of eGFP anchored on the SLB via click chemistry at different surface densities, corresponding to Figure 6. As the surface density of eGFP increases, the fluorescence recovery in the FRAP experiment becomes slower, implying the formation of protein condensates. The corresponding time courses of normalized fluorescence recovery at the bleached area are shown on the right. (Scale bar: 10 μm.)

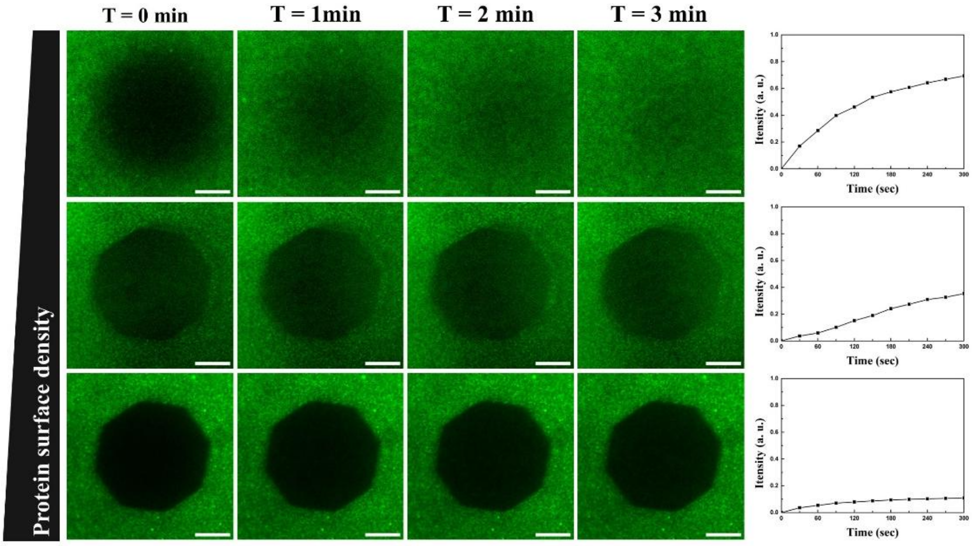


Figure S11. FRAP experiments of mNeonGreen anchored on the SLB via click chemistry at different surface densities (corresponding to Figure 6). As the surface density of mNeonGreen increases, the fluorescence recovery in the FRAP experiment shows a strong reduction in recovery rate, implying the formation of protein condensates. The corresponding time courses of normalized fluorescence recovery at the bleached area are shown on the right. (Scale bar: 10 μm.)

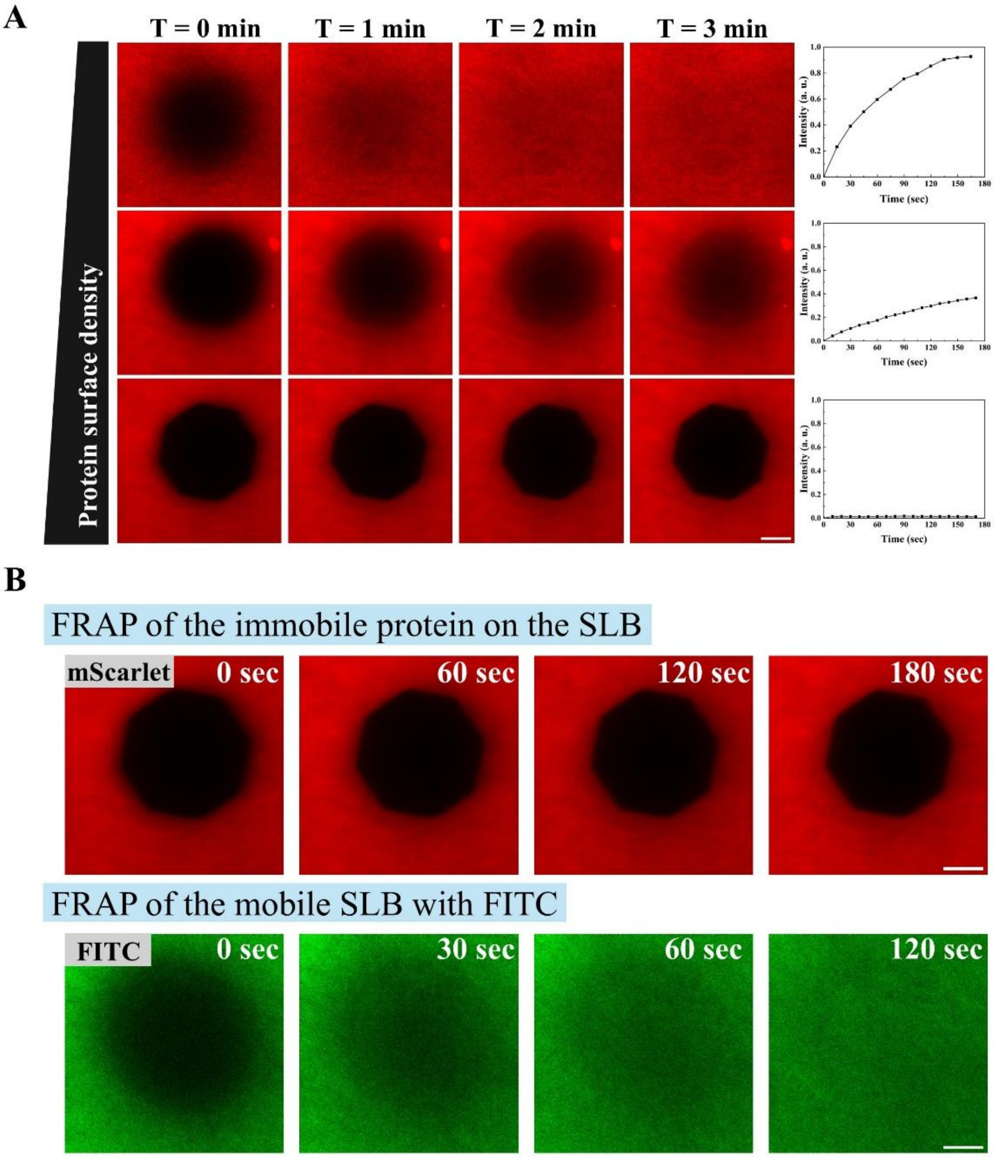


Figure S12. FRAP experiments of mScarlet anchored on the SLB via maleimide chemistry instead of click chemistry, corresponding to the data in Figure 7, at different surface densities (from 246 to 6084 molecules / μm²). As the surface density of mScarlet increases, the fluorescence recovery in the FRAP experiment shows a strong reduction in recovery rate, implying the formation of protein condensates. The corresponding time courses of normalized fluorescence recovery at the bleached area are shown on the right. These results indicate that the phase transition of the membrane-anchored protein driven by the crowding effect does not depend on the type of anchoring chemistry used. (B) The FRAP experiments of mScarlet from the third row in part A and the corresponding FITC lipid in the same SLB are compared. The fast fluorescence recovery of the FITC lipid on the membrane indicates that the membrane remains intact, while the protein forms immobile condensates on the membrane. (Scale bar = 10 μm.)

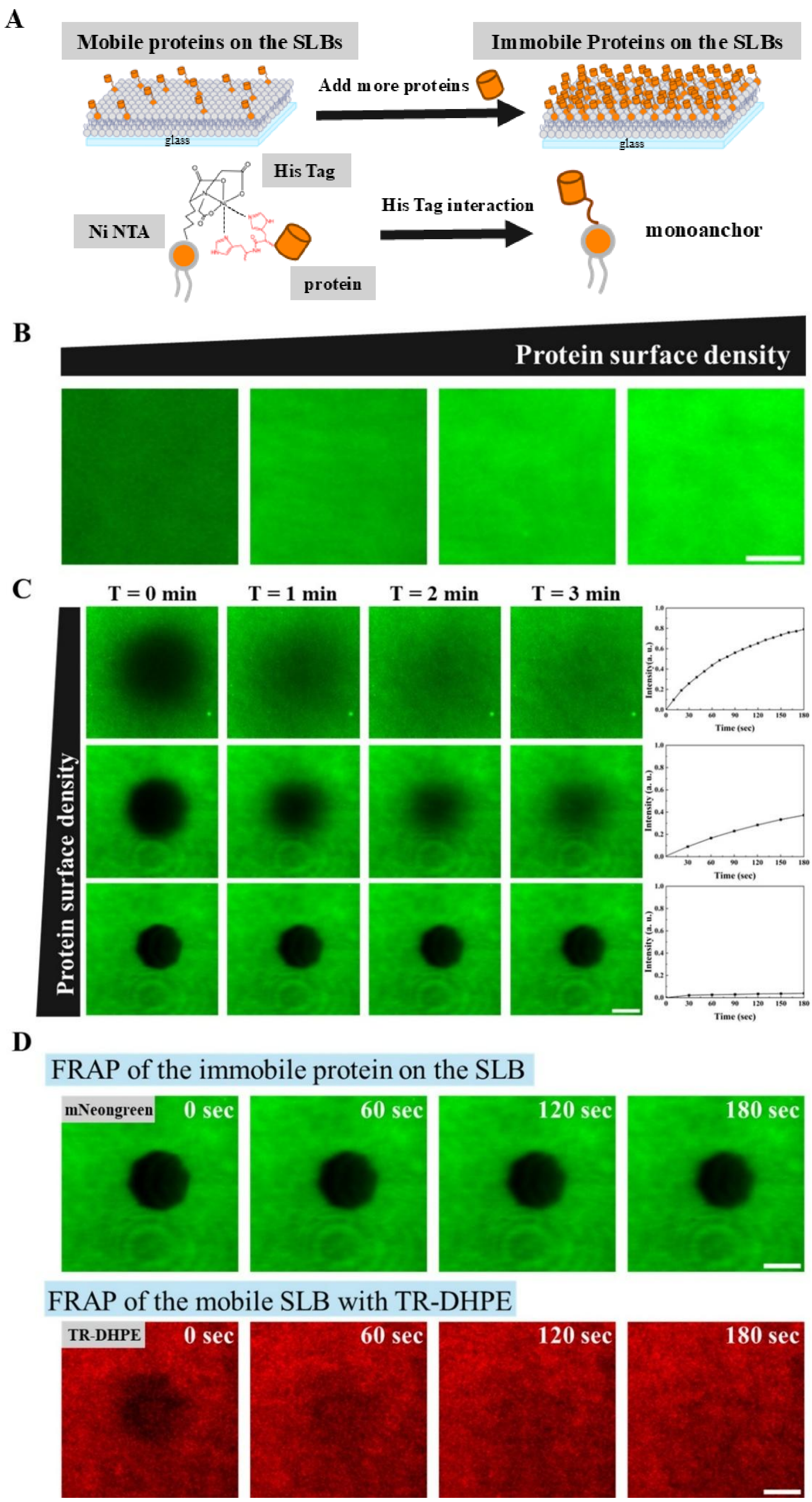
A
Mobile proteins on the SLBs
Immobile Proteins on the SLBs
Add more proteins
glass
glass
His Tag
Ni NTA
His Tag interaction
monoanchor
protein
B
Protein surface density
C
T = 0 min
T = 1 min
T = 2 min
T = 3 min
Protein surface density
Time (sec)
D
FRAP of the immobile protein on the SLB
mNeongreen
0 sec
60 sec
120 sec
180 sec
FRAP of the mobile SLB with TR-DHPE
TR-DHPE
0 sec
60 sec
120 sec
180 sec

Figure S13. mNeonGreen anchored on the supported lipid bilayers (SLBs) via His-tag chemistry. (A) The schematic illustrates how proteins are anchored to the SLB via His-tag chemistry. When the protein density increases, it leads to a crowding-induced phase transition. (B) Fluorescence images of mNeonGreen show that it is successfully anchored on the SLB via His-tag chemistry. By adjusting the concentration of mNeonGreen above the SLB, the surface density of mNeonGreen on the SLB can be controlled from 341 to 6118 molecules / $\mu m^2$. (C) FRAP experiments of mNeonGreen under different surface densities show that the fluorescence recovery rate exhibits a strong reduction as the surface density increases. The corresponding time courses of normalized fluorescence recovery at the bleached area are shown on the right. These results indicate the formation of immobile mNeonGreen protein condensates at high surface density, consistent with a crowding-induced condensation process. (D) The FRAP experiments of mNeonGreen and Texas Red–labeled lipids are compared. The fast fluorescence recovery of the Texas Red lipid on the membrane indicates that the membrane remains intact, while mNeonGreen forms immobile condensates on the membrane. (Scale bar = 10 μm.)
